# Biochemical and structural insights into SARS-CoV-2 polyprotein processing by Mpro

**DOI:** 10.1101/2022.05.27.493767

**Authors:** Ruchi Yadav, Valentine V. Courouble, Sanjay K. Dey, Jerry Joe E.K. Harrison, Jennifer Timm, Jesse B. Hopkins, Ryan L. Slack, Stefan G. Sarafianos, Francesc X. Ruiz, Patrick R. Griffin, Eddy Arnold

**Affiliations:** Center for Advanced Biotechnology and Medicine (CABM), Rutgers University, Piscataway, NJ, USA; Department of Chemistry and Chemical Biology, Rutgers University, Piscataway, NJ, USA; Department of Molecular Medicine, The Scripps Research Institute, Jupiter, FL, USA; Skaggs Graduate School of Chemical and Biological Sciences, The Scripps Research Institute, Jupiter, FL, USA; Department of Chemistry, University of Ghana, Legon, Ghana; BioCAT, Department of Physics, Illinois Institute of Technology, Chicago, IL, United States; Division of Laboratory of Biochemical Pharmacology, Division of Infectious Diseases, Department of Pediatrics, Emory University School of Medicine, Atlanta, GA, USA; Children’s Healthcare of Atlanta, Atlanta, GA, USA.; Department of Integrative Structural and Computational Biology, The Scripps Research Institute, Jupiter, FL, USA; Department of Molecular Medicine, UF Scripps Biomedical Research, University of Florida, Jupiter, FL, USA

## Abstract

SARS-CoV-2, a human coronavirus, is the causative agent of the COVID-19 pandemic. Its ∼30 kb RNA genome is translated into two large polyproteins subsequently cleaved by viral papain-like protease and main protease (Mpro/nsp5). Polyprotein processing is essential yet incompletely understood. We studied Mpro-mediated processing of the nsp7-10/11 polyprotein, whose mature products are cofactors of the viral replicase, identifying the order of cleavages as: 1) nsp9-10, 2) nsp8-9/nsp10-11, and 3) nsp7-8. Integrative modeling based on mass spectrometry (including hydrogen-deuterium exchange and cross-linking) and X-ray scattering yielded three-dimensional models of the nsp7-10/11 polyprotein. Our data suggest that the nsp7- 10/11 structure in complex with Mpro strongly resembles the unbound polyprotein, and that both polyprotein conformation and junction accessibility determine the preference and order of cleavages. Finally, we used limited proteolysis assays to characterize the effect of a series of inhibitors/binders on Mpro processing of nsp7-11 and Mpro inhibition using a polyprotein substrate.

**Teaser:** We elucidated the structural basis of order of cleavage of SARS-CoV-2 nsp7-11 polyprotein, with implications for Mpro inhibition.

## Introduction

The severe acute respiratory syndrome coronavirus 2 (SARS-CoV-2; CoV-2), a member of the *Coronaviridae* family, is responsible for the ongoing COVID-19 global pandemic (1). The toll of CoV-2 is extraordinary in terms of worldwide repercussions in the number of infected people, deaths and pace of infection spread (https://covid19.who.int/). SARS-CoV-2 has an ∼30 kb (+)-sense RNA genome, one of the largest known of any RNA virus, that encodes 13 open reading frames (ORFs), including replicase (ORF1a/ORF1b), spike (S), envelope (E), membrane (M), nucleocapsid (N), and seven other ORFs that encode accessory proteins (2). ORF1a and ORF1b are translated to produce two large polyproteins, pp1a and pp1ab. These polyproteins are subsequently cleaved into 16 non-structural proteins (nsps) by virally encoded proteases: the papain-like protease (PLpro, a domain of nsp3), which cleaves junctions from nsp1 to the nsp4 N- terminus, and the main protease (Mpro, nsp5, 3C-like protease, 3CLpro), which cleaves junctions from the nsp4 C-terminus to nsp16 (3). The “polyprotein strategy”—used by most RNA viruses and retroviruses—allows for: i) a more compact genome, ii) regulation of activity through a precise temporal (*i.e.* stage of viral cycle) and spatial (*i.e.* subcellular location) cleavage pattern, and iii) cleavage intermediates having distinct and critical roles from those of the cleaved products, as shown for alphaviruses, picornaviruses, and noroviruses (*4–6*). Hence, coordinated processing of polyproteins is vital for regulating the viral life cycle.

Different polyprotein intermediates derived from Mpro-mediated pp1a/1b processing have been detected in other CoVs, including mouse hepatitis virus (MHV) (7, 8) and alphacoronavirus human CoV 229E (HCoV-229E) (9). CoV-2 and MHV belong to the betacoronavirus genera with the latter being a good surrogate mouse model for studying CoV-2 infection and biology (*10–12*). Notably, mutations in the junction sites within the MHV nsp7-10 polyprotein were found to be lethal for viral replication, with the exception of the nsp9-10 site, where mutations led to a crippled mutant virus (8). Additionally, a polyprotein intermediate of ∼150 kDa corresponding to nsp4-10/11 has been detected in pulse-chase experiments (*13, 14*). Reverse genetic studies with temperature-sensitive mutants in MHV suggest that nsp4-11 could serve as a scaffold where replicative enzymes (nsp12, nsp14, nsp16) may dock to perform their activities on the viral RNA (*15*). Alternatively, they may indicate that mutation of this intermediate perturbs Mpro processing (*16*). Thus, the functional role(s) of nsp4-10/11 in virus replication remain unclear. Additionally, the subcellular localization of nsp7 to nsp10 has been studied for several CoVs using immunofluorescence microscopy and cryo-electron microscopy/tomography. pp1a/1ab are anchored to the endoplasmic reticulum (ER) membranes by flanking transmembrane domains (TMD) of nsp4 and nsp6, along with membrane-spanning nsp3. This topology results in the membrane-anchored Mpro being exposed to the cytosol along with most likely the nsp7-10/7-11 region (7, 17-20). Moreover, data from CoVs and other RNA viruses suggest that “convoluted membranes” (the precursors of the coronavirus replication organelles formed by double- membrane vesicles) may be the main site of viral gene expression and polyprotein processing. However, it should be noted that these labeling techniques cannot distinguish between mature nsps and polyprotein intermediates.

More recently in CoV-2 infected cells, the identification of viral cleavage sites at nsp4, nsp8-9, and nsp10-12 junctions at different post-infection time points has validated the presence of polyprotein intermediates and thus garnered support for further investigation into their functional relevance and structures (3). Krichel and co-authors have applied a structure-function approach to investigate the processing of the SARS-CoV nsp7-10 and MERS-CoV nsp7-11 polyproteins *in vitro* using native mass spectrometry (MS) (*21, 22*). Their results emphasized the critical role of the polyprotein conformation and the structural environment of the cleavage junctions in determining cleavage order, as the order of processing was previously inferred by determining the specific activity of Mpro cleavage on short oligopeptide sequences comprising the cleavage junctions (*23*).

One of the most investigated CoV-2 targets has been Mpro with ∼500 PDB structural depositions (https://rcsb.org/covid19). These structures include Mpro in both immature forms (*24*), as well as its mature apo form [(https://rcsb.org/covid19) (*25, 26*)]. Furthermore, there are multiple structures of Mpro bound with inhibitors (*27, 28*)—including the recently FDA- approved Pfizer inhibitor [PF-07321332, nirmatrelvir (NMTV)], (*29*)—and small molecules and fragment binders (*30–32*), and several structures with peptide substrates and products (*33–35*). Even with these efforts and with >2000 SARS-CoV-2 PDB depositions, no CoV polyprotein structures have been reported to date (https://rcsb.org/covid19). Indeed, despite their importance in the viral life cycle, polyprotein structural knowledge is very underrepresented in comparison to the multitude of solved structures of mature, post-cleavage proteins (6).

Herein, we have employed a multi-pronged approach to study the structural basis of processing of the CoV-2 nsp7-10 and nsp7-11 polyprotein(s) by Mpro *in vitro*, given their highly dynamic nature and multidomain organization. We have characterized the processing kinetics through gel-based and pulse-labeling MS techniques, as well as the footprint of the polyproteins on Mpro and vice versa. We also determined the integrative structures of the nsp7-11 and nsp7-8 polyproteins (by MS, small-angle X-ray scattering (SAXS), and molecular modeling). These experiments allowed us to rationalize the order of processing of the polyprotein by Mpro and provided insights into binding of the polyprotein substrate to Mpro. Finally, taking advantage of the vast number of Mpro-ligand structures, we identified a set of binders (with some displaying antiviral activity) overlapping with regions of Mpro relevant for polyprotein binding outside of its active site, and probed them in limited proteolysis inhibition assays including the full-length polyprotein substrate. Altogether the information gathered from this study improves our understanding of the role of polyproteins in SARS-CoV-2 viral replication.

## Results

### nsp7-10/11 polyprotein processing by Mpro using limited proteolysis

Polyprotein processing in *Coronaviruses* is a precise and tightly regulated process (8, 16, 36). We first expressed and purified the nsp7-11 and nsp7-8 polyproteins, and wild-type Mpro (**Fig. S1 in the Supplementary Materials, SM)**, to assess the proteolytic cleavage order of SARS-CoV-2 polyproteins. Next, we conducted a semi-quantitative proteolysis assay of the nsp7- 11 polyprotein with Mpro using SDS-PAGE as a readout **(Fig. 1A, S2)**. Analysis of the nsp7-11 polyprotein processing revealed that the nsp9-10 junction was cleaved initially (starting ∼30 min), followed by simultaneous cleavage of the nsp8-9 and nsp10-11 junctions (starting ∼2 h), and finally the nsp7-8 junction (starting ∼4 h) **(Fig. 1A** and **1B**). This order of cleavage is identical to the polyprotein processing order reported for SARS-CoV (CoV-1), which was expected given their high amino acid sequence conservation (*22*). Analysis of the nsp7-10 polyprotein processing yielded similar results **(Fig. S3).** This suggests that the presence of nsp11 does not affect the polyprotein cleavage order. Moreover, altering ratios of Mpro to polyprotein had no effect on the cleavage order (1:2 and 1:12 molar ratio for nsp7-10 and nsp7-11 limited proteolysis, respectively), further supporting the specificity of Mpro and the lack of a concentration-dependent cleavage effect. Interestingly, after 24 h of exposure to Mpro, the nsp7-8 junction was not completely cleaved. The limited proteolysis assay with the nsp7-8 intermediate polyprotein showed that nsp7-8 was also not fully cleaved after 24 h (1:10 molar ratio of Mpro:nsp7-8) **(Fig. S4),** suggesting that the structural environment around the nsp7-8 junction impedes efficient Mpro cleavage with respect to the other junction sites.

**Fig. 1.**
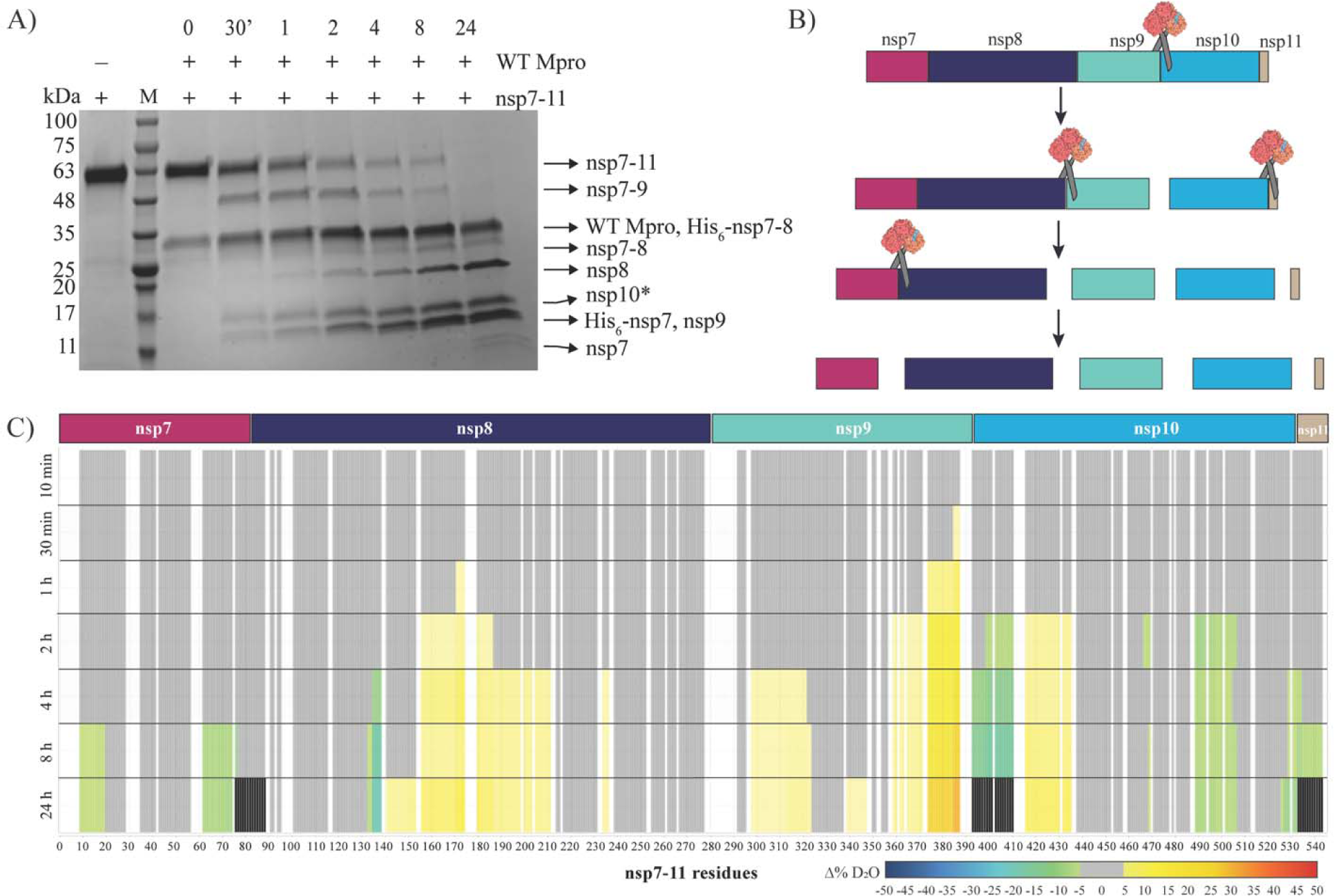
*In vitro* analysis of the nsp7-11 polyprotein processing by WT Mpro. (A) SDS-PAGE showing the limited proteolysis of nsp7-11 polyprotein by Mpro over a time course of 24 h. +/- shows the presence or the absence of the respective proteins. The lane labeled as M is the protein marker. Black arrows on the right indicate the proteins generated from the cleavage of nsp7-11 polyprotein by Mpro. (B) Schematic representation of the cleavage order of the nsp7-11 polyprotein by Mpro. (C) Pulsed HDX-MS analysis of nsp7-11 with Mpro. Color scale represents changes in deuterium uptake over the course of the cleavage reaction, with gray representing no significant change in deuterium uptake, white denoting no sequence coverage, and black representing residues within peptides that are no longer identifiable.

### Pulsed HDX-MS reveals cleavage order of polyprotein by Mpro

To gain further structural insight into the polyprotein processing with Mpro, we conducted pulsed HDX-MS. Briefly, we incubated nsp7-11 with Mpro at an equimolar ratio and the cleavage reaction was allowed to proceed over 24 h. Aliquots of the reaction were taken at various time intervals and incubated in deuterated buffer for 30 s before being quenched, flash-frozen, and stored until ready for MS analysis. All time points were compared to nsp7-11 without Mpro to observe changes in solvent exchange occurring in the polyprotein over the course of the proteolytic process **(Fig. 1C, S5, Table S1)**.

We observed increased deuterium exchange (destabilization/decreased hydrogen bonding) in the nsp9 C-terminal region at 30 min **(Fig. 1C)**, indicative of increased conformational mobility in this region of the polyprotein upon interaction with Mpro. Deuterium exchange continued to increase over time in this region as compared to the protein in the absence of enzyme. This change was followed by increased deuterium exchange of the nsp10 N-terminal region, indicating increased conformational mobility in this region. Concomitantly, we observed decreased solvent exchange at the cleavage junction residues, suggesting reduced solvent accessibility due to Mpro binding. Over the time course, we also observed a decrease in signal intensity of the nsp9-10 junction site spanning peptides indicative of cleavage. At 24 h we were no longer able to detect these peptides in the mass spectrometer indicating full cleavage of the nsp9-10 site **(Fig. S5)**. These observations suggest that the nsp9-10 junction is being cleaved first, consistent with our SDS-PAGE analysis **(Fig. 1A)**.

The nsp8-9 and nsp10-11 junctions appeared to be simultaneously cleaved next as both junctions showed changes in deuterium exchange starting at 4 h. At the nsp9 N-terminal region, we observed increased deuterium exchange compared to the protein in the absence of the enzyme, indicating increased solvent exchange in this region. As we were unable to detect peptides specifically spanning the nsp8-9 junction, we could not determine the exact timing of full cleavage. Meanwhile, at the nsp10-11 junction site, we observed decreased solvent exchange and reduced signal intensity in the peptides spanning the junction site until 24 h where they were no longer identified in the mass spectrometer to suggest full cleavage. While it appears that nsp11 is no longer identified at 24 h, this is due to our inability to detect any nsp11 only peptides, as it is only 13 amino acids in length, such that all the peptides covering nsp11 are also covering the junctions **(Fig. S5)**. Nevertheless, it was clear that nsp8-9 and nsp10-11 are cleaved simultaneously, following cleavage at the nsp9-10 site.

The nsp7-8 site is cleaved last, as we did not observe any changes in deuterium uptake near the nsp7-8 junction until 8 h. While we do not observe decreased solvent exchange in the peptides spanning the nsp7-8 junction to indicate Mpro binding at this site, we did not detect these peptides in the mass spectrometer at 24 h suggesting that the nsp7-8 junction is cleaved by Mpro.

Interestingly, we also observed changes in solvent exchange away from the junction sites in nsp7 and nsp8. The protection from exchange (stabilization/increased hydrogen bonding) within both nsp7 and nsp8, suggests that nsp7 and nsp8 associate into a heterodimer after their release from the polyprotein. The pattern of protection observed in nsp7 agreed well with protection we observed in our prior differential HDX-MS analysis of the nsp7:nsp8 heterotetrameric complex (*37*). Unexpectedly, increased exchange (destabilization) within nsp8 was also observed starting at 2 h. These peptides in nsp8 that showed increased exchange demonstrated EX1 behavior as revealed by detection of two distinct deuterated ion distributions (bimodal) for the same peptide (*38*). Under native conditions this behavior has been shown to be a result of multiple intermediate conformational protein states (*39–42*). The observed EX1 behavior in nsp8 could be explained by either increased flexibility of the nsp8 N-terminus adopting multiple conformations—previously documented (*21, 37*)—and/or by the simultaneous presence of both mature nsp8 protein and nsp7-8 polyprotein within the samples.

Additionally, nsp10 also showed decreased deuterium uptake away from the junction site. Comparing the solvent exchange profile of nsp7-11, nsp7-10, and individual nsp10 **(Fig. 3A)**, we observed that nsp10 has the greatest deuterium uptake in nsp7-11 while nsp7-10 and nsp10 showed similar solvent exchange profiles, suggesting that the presence of nsp11 destabilizes nsp10. Specifically, the regions of protection from solvent exchange observed in nsp10 during the pulsed HDX-MS experiment align with the residues showing decreased deuterium uptake in mature nsp10 and nsp7-10. Moreover, the pulsed HDX-MS with nsp7-10 **(Fig. S6)** did not show any changes in deuterium exchange in nsp10, as expected from the comparison of the solvent exchange profiles of nsp7-10 and mature nsp10. This confirms that released nsp10 does not interact with other liberated proteins in solution and the observed decreased solvent exchange in nsp10 is due to the release of nsp11 from nsp10.

Overall, the pulsed HDX-MS results were consistent with the SDS-PAGE proteolytic results, showing the processing order to be: 1) nsp9-10, 2) nsp8-9 and nsp10-11, and finally 3) nsp7-8 **(Fig. 1B)**. Moreover, pulsed HDX-MS with nsp7-10 displayed similar results with the same processing order as well as interaction of mature nsp7 and nsp8 after their release **(Fig. S6)**.

### Differential HDX-MS demonstrates localized sites of interaction of C145A Mpro to polyprotein junction sites

Next, we used HDX-MS and XL-MS as complementary techniques to better understand the solution phase dynamics of the complex. Using differential HDX-MS, we compared nsp7-11 versus nsp7-11 in complex with C145A Mpro at an equimolar ratio **(Fig. 2A, S7A, Table S1).** Increased protection from solvent exchange was observed at all junction sites except the nsp10-11 junction. The nsp9-10 junction had the largest magnitude in reduction of solvent exchange which may suggest it to be the primary binding site on the polyprotein. This observation is consistent with the proteolysis SDS-PAGE **(Fig. 1A)** and pulsed HDX-MS results **(Fig. 1C)** that indicate the nsp9-10 junction to be the initial target of Mpro. Minimal alteration of solvent exchange was observed within the nsp subdomains outside of the junction sites, suggesting that Mpro interaction with the polyprotein is favored at the junction site sequences, and that binding of Mpro does not induce significant long-range conformational changes in the polyprotein. Only nsp8 showed additional regions of protection from solvent exchange away from the junctions, specifically residues T120-M140 and K182-L213. These regions of the polyprotein are inherently more dynamic, as determined by higher intrinsic deuterium exchange **(Fig. 3A)**, and thus the observed protection suggests that interaction with C145A Mpro is stabilizing the nsp8 N-terminal region.

**Fig 2.**
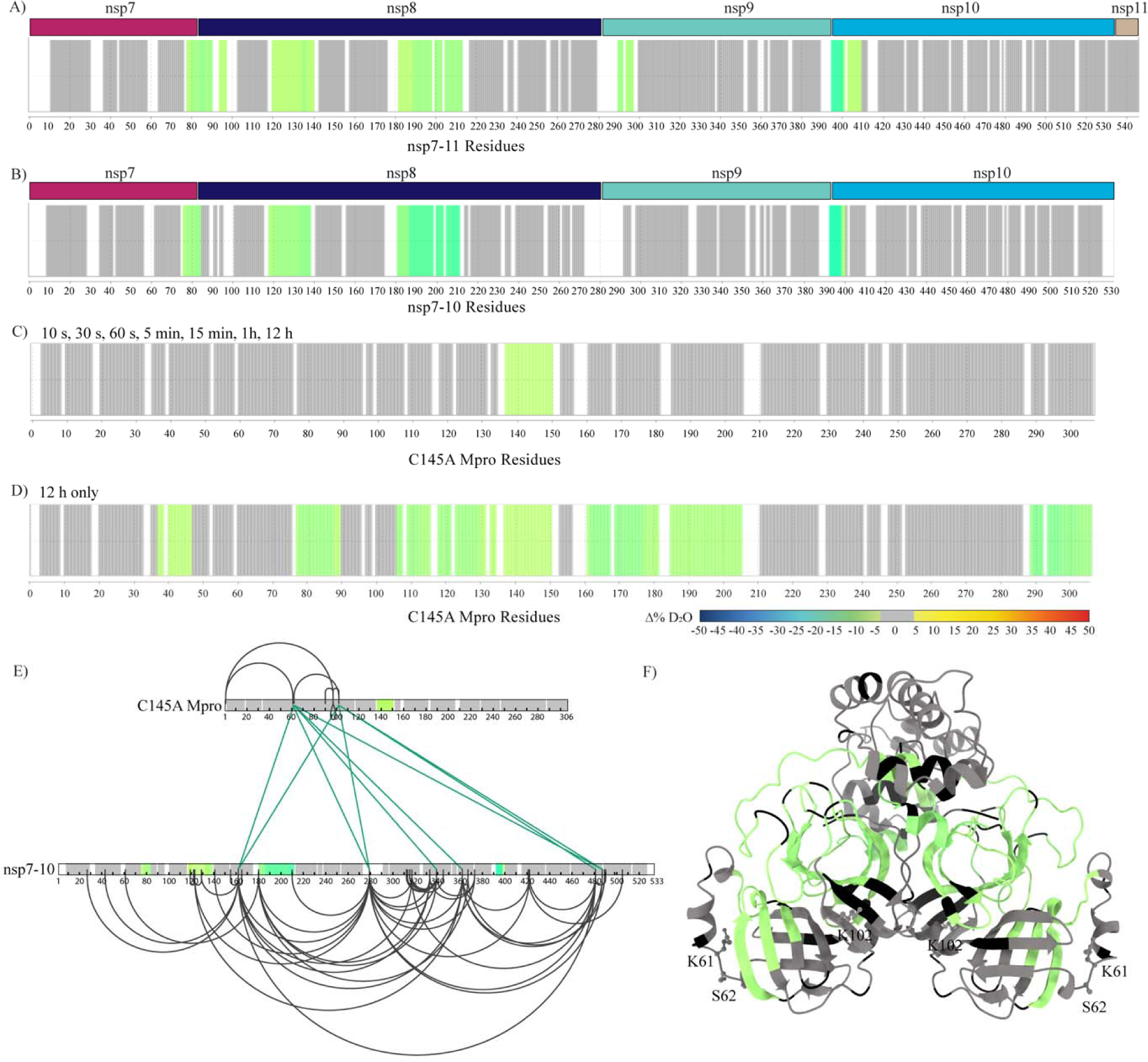
HDX-MS and XL-MS reveal the in-solution dynamics of the Mpro polyprotein complex. Consolidated differential HDX-MS results of (A) nsp7-11 vs nsp7-11 in complex with C145A Mpro, (B) nsp7-10 vs nsp7-10 in complex with C145A Mpro, (C) C145A Mpro vs C145A Mpro with nsp7-11 over time course up to 1 h and including 12 h, and (D) C145A Mpro vs C145A Mpro with nsp7-11 at 12 h only. All consolidated differential HDX-MS results are colored based on change in percent deuterium as described in scale bar, with regions showing no significant change in deuterium in gray and regions with no sequence coverage in white. (E) Overlay of HDX-MS and XL-MS results on nsp7-10 and C145A Mpro sequences. Observed intra Mpro and intra nsp7-10 crosslinks colored in black and inter C145A Mpro to nsp7-10 crosslinks colored in green. Consolidated changes in percent deuterium uptake are taken from **Fig. 2B and 2C**. (F) Overlay of HDX-MS results on C145A Mpro (modeled based on PDB 7DVY). C145A Mpro residues forming inter protein crosslinks with nsp7-10 are shown as sticks and labeled. Consolidated changes in percent deuterium uptake are taken from **Fig. 2D**.

Similar results were observed with the nsp7-10 polyprotein **(Fig. 2B, S7B, Table S1).** All the cleavage sites in nsp7-10 were protected from solvent exchange upon interaction with C145A Mpro. Additionally, the nsp9-10 junction showed greatest protection from solvent exchange, while only nsp8 showed additional regions of protection outside the junctions. These results indicate that nsp11 does not alter the polyprotein interactions with C145A Mpro.

### Differential HDX-MS validates polyprotein binding to C145A Mpro beyond the active site pocket

Next, we profiled changes in solvent exchange of C145A Mpro bound to nsp7-11 at an equimolar ratio but did not observe any significant protection from solvent exchange in case of C145A Mpro (data not shown). However, C145A Mpro deuterium uptake was relatively low over the experimental time course up to one hour, suggesting that the Mpro dimer is very stable. To increase the observable window to detect protection from solvent exchange, we incubated C145A Mpro and the C145A Mpro:nsp7-11 complex in deuterium buffer for 12 h. Overall, deuterium uptake was increased over this longer time course and significant protection from solvent exchange was observed in the active site region containing the C145A mutation only when in the presence of polyprotein **(Fig. 2C, S7C, Table S1).** Additional regions of the enzyme were shown to be protected from solvent exchange at 12 h incubation in deuterated buffer only, including peptides spanning residues T45-S46, V77-L89, I106-F150, Y161-L205, and D289-Q306 **(Fig. 2D, S7D, Table S1).** These regions were mapped from the back side of the C145 Mpro active site pocket to the vicinity of the dimerization interface, hinting to a putative binding track for the rest of the nsp7-11 polyprotein outside of the nsp9-10 junction (see below for more details).

### XL-MS demonstrates additional contact sites between C145A Mpro and polyproteins

While the differential HDX-MS analysis of the C145A Mpro polyprotein complex reported on changes in protein backbone dynamics, we also analyzed the complex using XL-MS to probe protein side chain residency and reactivity. A total of nine inter-protein crosslinks were identified between C145A Mpro and nsp7-10 **(Fig. 2E)**. The three C145A Mpro residues (K61, S62, and K102) that form inter-protein crosslinks with nsp7-10 were mapped to the catalytic domain **(Fig. 2F)**. When these crosslinks were mapped alongside the HDX-MS data, we observed that the inter-protein crosslinks map to residues outside of regions showing protection from solvent exchange **(Fig. 2E)**. Accordingly, these crosslinks represent additional contact sites between the polyprotein and Mpro that may help stabilize the complex to position the junctions into the active site. For example, K162 within nsp8 is located between the two regions of nsp8 showing protection from solvent exchange **(Fig. 2B)** and forms an interprotein crosslink with S62 and K102 in Mpro. This further supports that interaction with C145A Mpro stabilizes nsp8 conformation.

### HDX-MS profile of polyprotein revealed similar secondary structure elements to individual nsps

The HDX-MS intrinsic exchange profiles of nsp7-11, nsp7-10, and nsp7-8 polyproteins all revealed similar solvent exchange behavior **(Fig. 3A)**, which suggests that all polyproteins share similar secondary structure elements and overall conformation. Additionally, the HDX-MS intrinsic exchange profiles of the polyproteins largely resembled the intrinsic exchange profiles of individual nsp7, nsp8, and nsp9 **(Fig. 3A)**. This suggests that the secondary structures within the polyproteins remain largely unchanged in these mature nsps. The only exception is mature nsp10, which by itself or within nsp7-10, shows reduced deuterium uptake compared to nsp10 within the nsp7-11 polyprotein. This suggests that nsp10 is destabilized (decreased hydrogen bonding) when bound to nsp11.

**Fig 3.**
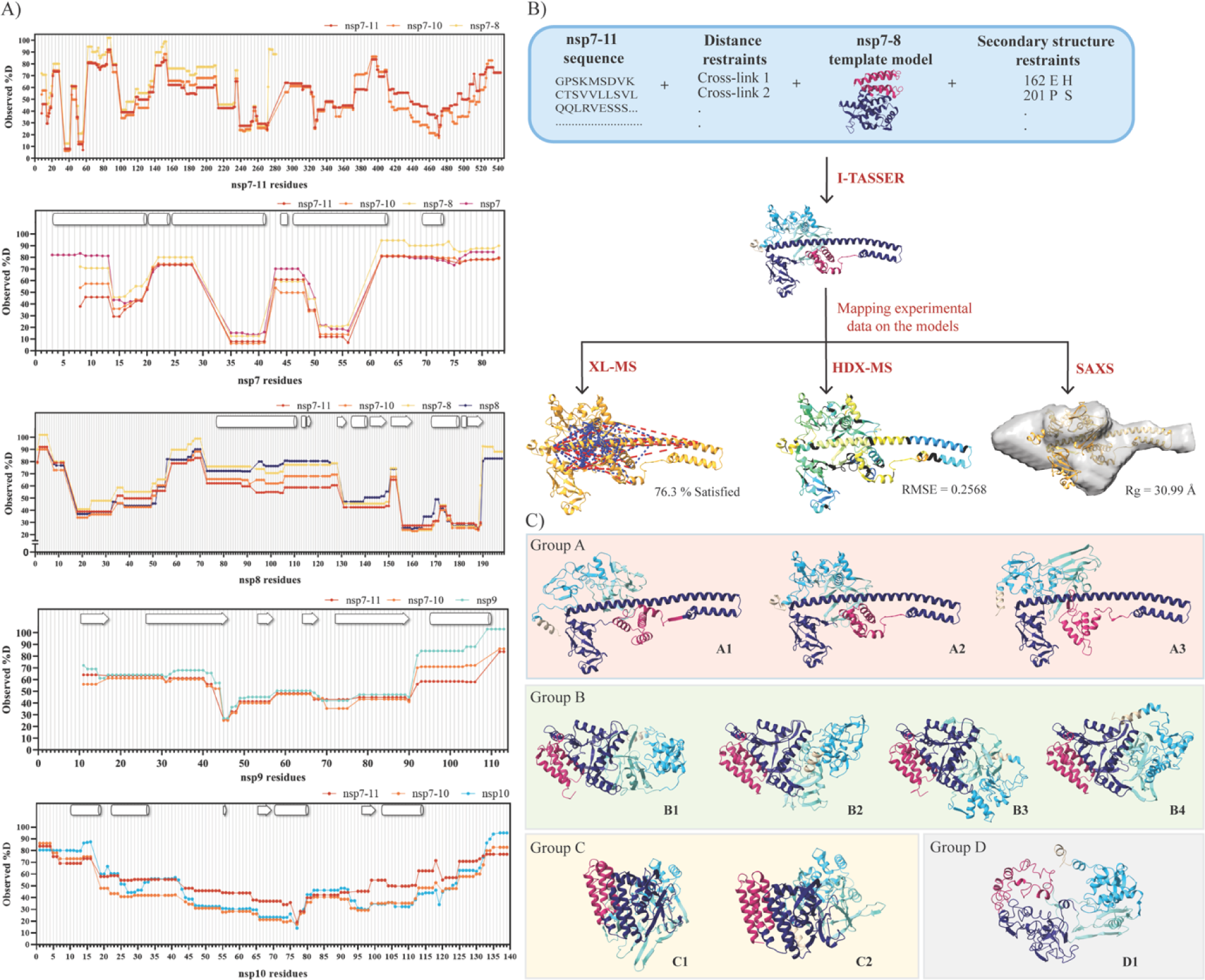
Integrative structural modeling generates an ensemble of nsp7-11 models. (A) Plots of observed percent deuterium per residue for nsp7, nsp8, nsp9, nsp7-8, nsp7-10, and nsp7-11. Secondary structures from PDB 6YHU for nsp7 and nsp8, PDB 6WXD for nsp9, and PDB 6ZPE for nsp10 are drawn on plots with α-helices shown as barrels, L-strands shown as arrows and coils shown as rectangles. (B) Scheme of integrative structural modeling workflow for the nsp7- 11 polyprotein. One model is shown to represent all ten generated. (C) Top ten nsp7-11 models grouped into four representative tertiary structures. Models are colored by nsp with nsp7 in magenta, nsp8 in purple, nsp9 in teal, nsp10 in cyan, and nsp11 in tan.

### Integrative structural modeling of the nsp7-11 polyprotein structure proposes an ensemble of different conformations

Next, we turned to an integrative structural modeling approach using multiple experimental techniques to account for biases inherent to each technique. We used the *ab initio*- based I-TASSER algorithm that allows incorporation of experimental restraints (*43*). We decided to focus the integrative modeling efforts on nsp7-11 as translation of ORF1a includes nsp11.

We first used analytical size exclusion chromatography (SEC) coupled to multi-angle light scattering (MALS) and SAXS detection (SEC-MALS-SAXS) to analyze the assembly state and structural features of the polyprotein in-solution. The SEC of the nsp7-11 polyprotein showed two peaks suggesting the presence of two different states: monomer and dimer, with the monomeric form being predominant **(Fig. S8A)**. The MALS analysis was used to calculate the molecular weights of the two identified peaks for nsp7-11 (∼60 and ∼110 kDa). SAXS analysis was conducted for both oligomeric states to understand the arrangement of these polyproteins in solution. Concretely, evolving factor analysis was used for separating the scattering of the monomer and dimer components **(Fig. S8B)** in a model-independent way (*44*). Both states yielded a linear Guinier plot indicating the presence of stable protein sample with no aggregation **(Fig. S8C)**. The bell-shaped (Gaussian) curve at lower q values in the Kratky plot showed that the sample contains folded domains with no significant disorder **(Fig. S8D)**. The pair-distance distribution function, P(r), which is related to the shape of the sample, indicated a globular shaped protein for both the monomeric and dimeric forms of nsp7-11 **(Fig. S8E)**. The Rg and the Dmax values calculated from the P(r) are: 48.2 Å and 191 Å for the dimer, and 35.8 Å and 156Å for the monomeric nsp7-11 **(Table S2)**.

Subsequently, we applied the integrative structural modeling approach to predict the structure(s) of the monomeric state of the nsp7-11 polyprotein **(Fig. 3B)**, which may likely be the form binding to Mpro (based on the HDX-MS data, **see Fig. 2A, S7A**). Models were generated based on the amino acid sequence and the following experimental parameters: i) distance constraints from XL-MS, ii) secondary structure restraints from solved X-ray crystal structures of the mature nsp7 to nsp10 (guided by solvent exchange profiles from HDX-MS indicating similar secondary structures of nsps within the polyprotein or upon cleavage), and iii) various nsp7-8 polyprotein models **(Fig. S10A-F, more details in SM)**. A final ensemble of ten nsp7-11 models were binned into four representative conformational groups, which were all assessed using our gamut of techniques **(Fig. 3B-C)**.

When we compared the four model groups, the nsp7 helical bundle stood as the most conserved structural element in all models, except for Group D. The other nsp domains presented more diversity in structural conformations and orientations. Group A is defined by an extended helical N-terminus, with a “golf-club” conformation, observed in the CoV-1 nsp7:nsp8 complex (*45*) and in CoV-2 structures of nsp8 interacting with nsp7 and nsp12 (*46–48*). Groups B and C exhibit a more compact organization of nsp8, with Group B having nsp7, nsp8, nsp9, and nsp10 arranged linearly, while Group C has all domains arranged in a packed “sphere” and Group D presents an “open” conformation.

Despite these conformational differences, all the models satisfied most of the crosslinks, with distances equal to or less than 30 Å (upper limit distance for DSSO crosslinks) **(Fig. 4A, S11, Table S3)**. Specifically, Group B satisfied the greatest percentage of crosslinks while Group A and D had the highest number of violations. These violations mostly stemmed from the extended nsp8 N-terminal helix in Group A and the less ordered conformation in Group D, which suggests that the nsp7-11 polyprotein, and especially the nsp8 segment, samples multiple conformations in solution. This conclusion is also supported by the HDX-MS data, showing that the central region of the nsp8 N-terminal subdomain exhibited greater solvent exchange (higher percent deuterium) suggesting increased inherent dynamics **(Fig. S12D)**.

**Fig 4.**
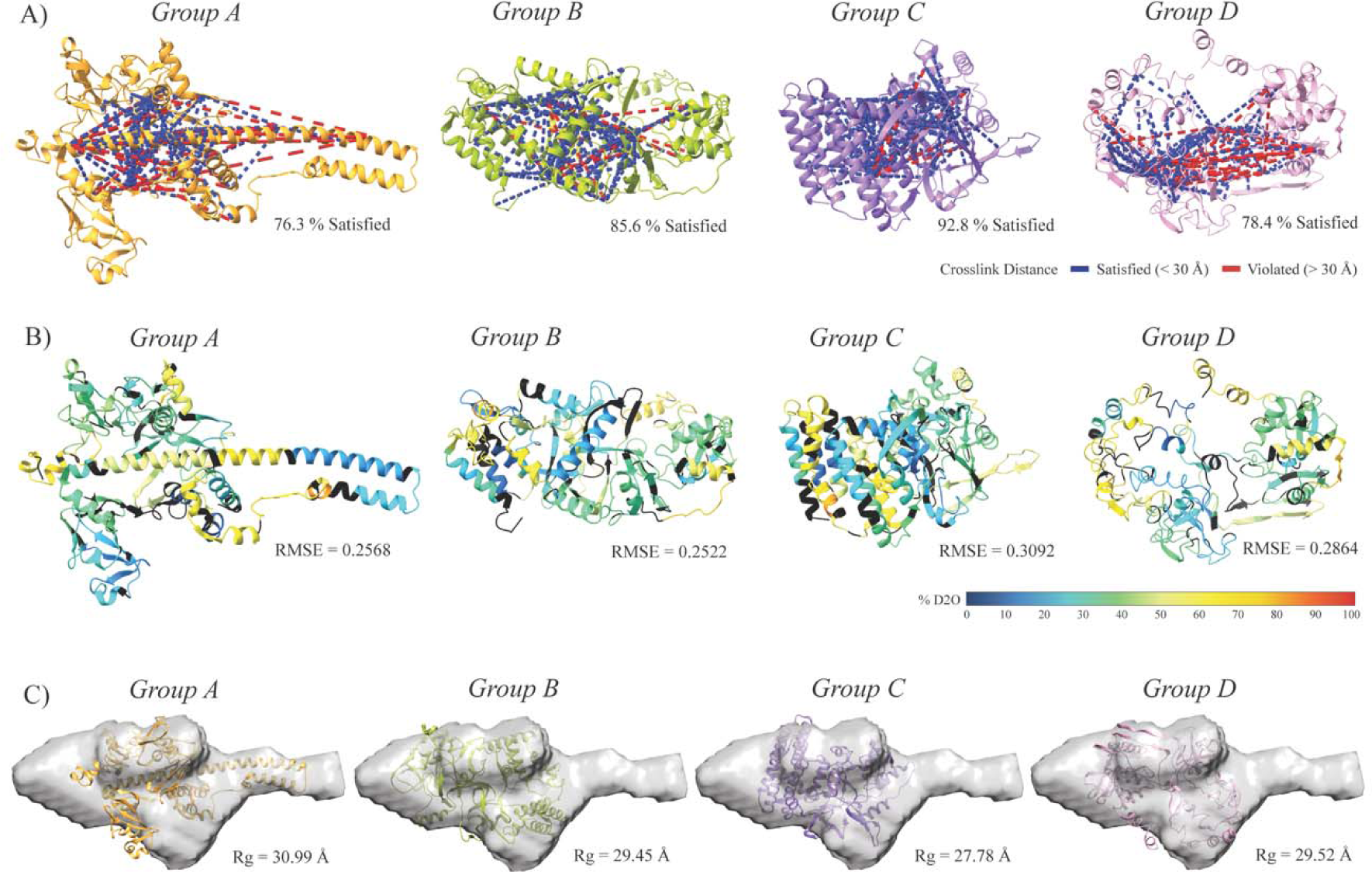
Assessment of representative nsp7-11 models based on experimental data. (A) Mapping of nsp7-11 intra-protein crosslinks onto representative nsp7-11 models. Satisfied crosslinks equal to or less than 30 L are shown in blue and violated crosslinks greater than 30 L are shown in red. Percent of crosslinks satisfied is reported under structure. (B) Representative nsp7-11 models are colored based on 10 s percent deuterium value. Black indicates no sequence coverage in the HDX-MS experiment. Agreement of model with experimental data as calculated by HDXer is reported as the RMSE under the model. (C) Fitting of representative nsp7-11 models into the SAXS envelope and Rg values are reported under the model.

Next, we evaluated the models with the HDXer software using the percent deuterium uptake values from our HDX-MS experiments as input (*49, 50*) **(Fig. 4B, S12A-B, Table S3)**. HDXer calculates deuterated fractions for peptide segments corresponding to the experimental data as a function of the experimental deuterium exposure times. We then plotted the computationally derived percent deuterium value at 10 s incubation in deuterated buffer for each model versus the experimentally determined percent deuterium value at 10 s incubation in deuterated buffer and calculated their Root Mean Square Error (RMSE). Group A demonstrated the lowest RMSE (best agreement), compared to other models. Group C and D models had the highest RMSE suggesting these conformations to be representative of models with too much rigidity or too much flexibility, respectively. Moreover, it was noted that nsp11 had the worst agreement with experimental HDX-MS data, while nsp7 had the best agreement **(Fig. S12C)**, which is likely due to the lack of nsp11 structures and abundance of X-ray crystal structures of nsp7 in complex with nsp8.

Next, a three-dimensional shape (bead model) was reconstructed for the monomeric nsp7- 11 from the scattering profile using DAMMIF/N from the ATSAS package (*51, 52*). As the SAXS scattering profile represents averaged scattering from all different possible orientations, it may be possible that many different shapes/orientations can generate the same scattering profile and therefore, for certain shapes it can be difficult to generate a bead model that correctly represents the solution shape. To assess whether a bead model uniquely fits the scattering data or if multiple models can fit the data, certain criteria are checked including ambiguity score, normalized spatial discrepancy (NSD) value, number of clusters, as well as parameters such as Rg, Dmax, and molecular weight (M.W.) values. Ambiguity score or “a-score” is the initial screening which informs about the number of possible shapes representing the same scattering profile. An “a-score” below 1.5 is usually indicative of a unique *ab initio* shape determination. In our case, 0.85 “a-score” suggested a unique 3D reconstruction. The Rg and Dmax obtained from the reconstructed model were close to those calculated from the P(r) function. The M.W. of the refined model was also comparable to the expected M.W. **(Table S2)**. Another important criterion to consider is NSD, which is used to evaluate the stability of the reconstruction. An NSD value less than 1.0 suggests fair stability of the reconstructions. DAMAVER reported 0.95 NSD for our reconstruction, which is on the borderline of a stable reconstruction. DAMCLUST created nine different clusters, suggesting that several different shapes in solution could have generated the same scattering profile. While the ambiguity score and comparable Rg, Dmax and M.W. values favored the bead model reconstruction, other criteria such as NSD and number of clusters suggested heterogeneity in the reconstruction. The higher NSD value and multiple clusters are likely due to nsp7-11 adopting multiple conformations. As stated earlier, the central segment of nsp8 is highly flexible and dynamic as suggested by greater solvent exchange in HDX, which could lead to heterogeneity in the conformations. Out of the four representative conformational groups, Group A showed the best fit with the reconstructed SAXS envelope, as the extended nsp8 helix fit into the elongated extension of the envelope. Interestingly, the less ordered and open conformation of the Group D model appeared to fit better in the SAXS envelope compared to the more ordered and compact structure of Group C models. We also compared the calculated scattering profile for the models to the experimental scattering profile. The χ2 and Rg values for Group A showed the greatest agreement with experimental data **(Fig. 4C, Fig. S13, Table S3)**.

In summary, the assessment of the ten models using HDX-MS, XL-MS, and SAXS highlighted that nsp7-11 can sample four major conformations **(Table S3)**. To note, our integrative structure modeling approach cannot ascertain the abundance of the different conformers within the ensemble. Group A conformers adopted an extended nsp8 helix with good agreement with HDX-MS and SAXS data but poor agreement with XL-MS. Group B conformers showed linear nsp organization with good XL-MS agreement but average agreement with HDX-MS and poor SAXS agreement. Group C conformers were arranged as a packed spherical structure with poor HDX-MS and SAXS agreement but good XL-MS agreement. Finally, Group D conformers had the most dynamic conformations (i.e., fewer ordered secondary structural elements) and showed good agreement to SAXS data but average agreement to HDX-MS and poor XL-MS agreement **(Table S3)**.

### The ensemble of nsp7-11 models unveils the interplay between cleavage junction conformation and accessibility to determine preference and order of cleavage

Next, we evaluated the structural environment of the cleavage junctions in the ensemble of nsp7-11 models to understand the influence of polyprotein substrate conformation and accessibility in processing **(Fig. 5, S14, Table S3)**. In general, all the junctions (except for nsp8-9, which was just partially covered by HDX-MS) (**Fig. 2A**) showed high levels of solvent exchange (high percent deuterium values), consistent with the fact that the cleavage regions should be accessible for proteolysis to occur. The combination of secondary structure and accessible surface area for Groups B and C was most consistent with the processing order we determined by limited proteolysis and pulsed HDX-MS **(Fig. 5, S14, Table S3)**. Comparing all the junctions, the nsp9- 10 junction, which was the first to be cleaved, was the most exposed junction in all the models and adopted a random coil in all but one model, which may best facilitate interaction with Mpro. On the other hand, the nsp7-8 junction, which was the last to be cleaved, was more hindered and mostly adopted an α-helical conformation, which may entail a slow cleavage event. Interestingly, for Group A models, nsp7-8 junction appeared to be the most accessible junction, which ultimately lends to our conclusion that the polyprotein is likely sampling multiple conformations with some being more amenable to proteolytic processing than others.

**Fig 5.**
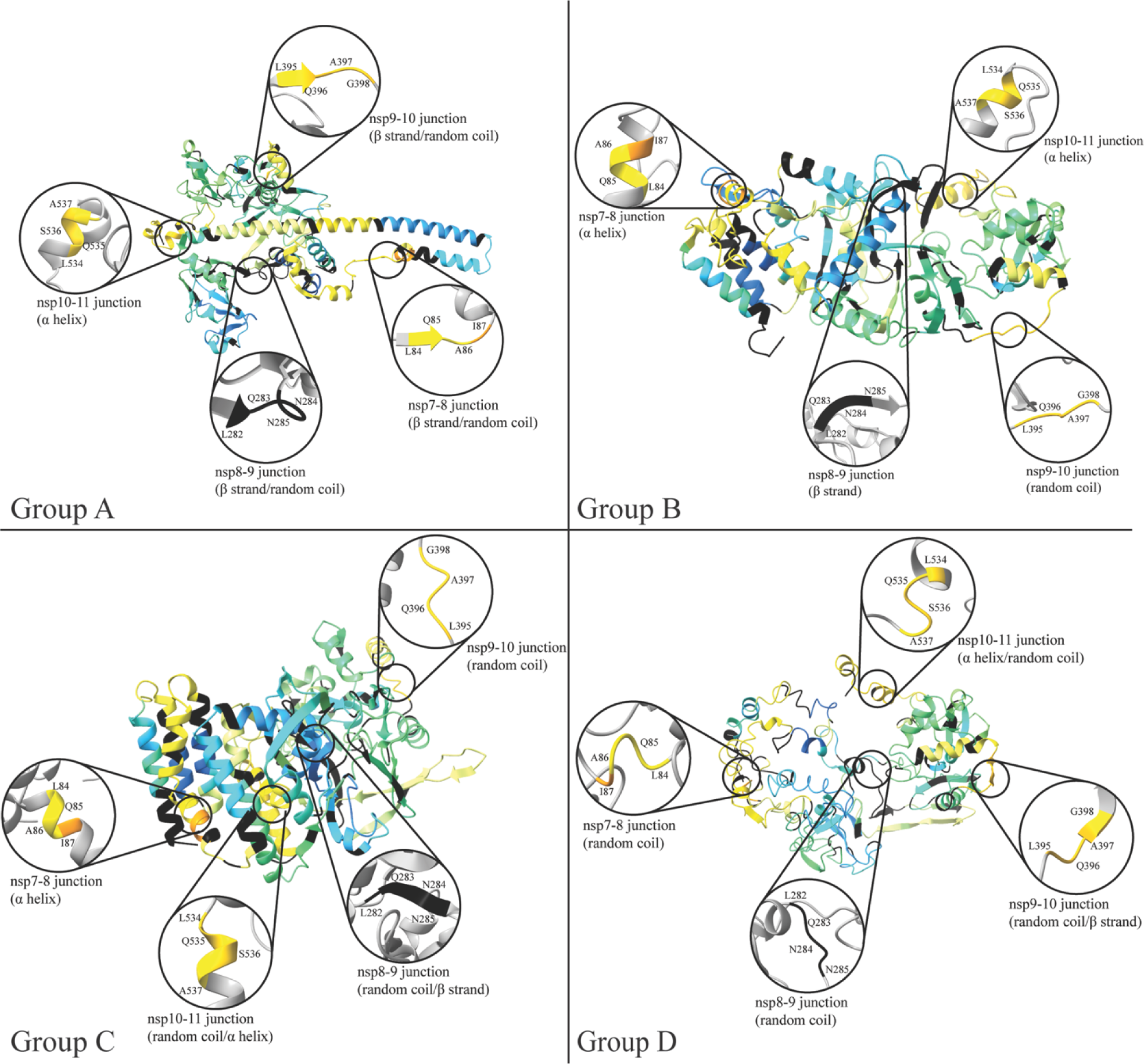
Assessment of junction site in representative nsp7-11 models. Analysis of secondary structure elements of junction sites in representative nsp7-11 models. Models are overlaid with 10 s deuterium uptake values from **Fig. 4B**.

### Probing nsp7-11 binding to Mpro with small molecule binders

To further understand the implications of polyprotein binding to Mpro outside its active site—studied first via HDX- and XL-MS—regarding proteolytic processing, we leveraged the limited proteolysis assay using the nsp7-11 polyprotein as Mpro substrate to measure inhibition by active site and non-active site binders of Mpro identified through crystallography (*30, 32*). Specifically, we selected small molecule binders—some of them presenting antiviral activity, but most of them not tested in enzymatic assays (*32*)—overlapping with the Mpro regions showing protection in the differential HDX-MS of nsp7-11 on C145A Mpro at 12 h **(Fig. 6A and Table S4)**. We used the FDA-approved drug nirmatrelvir (NMTV) as a positive control.

**Fig 6.**
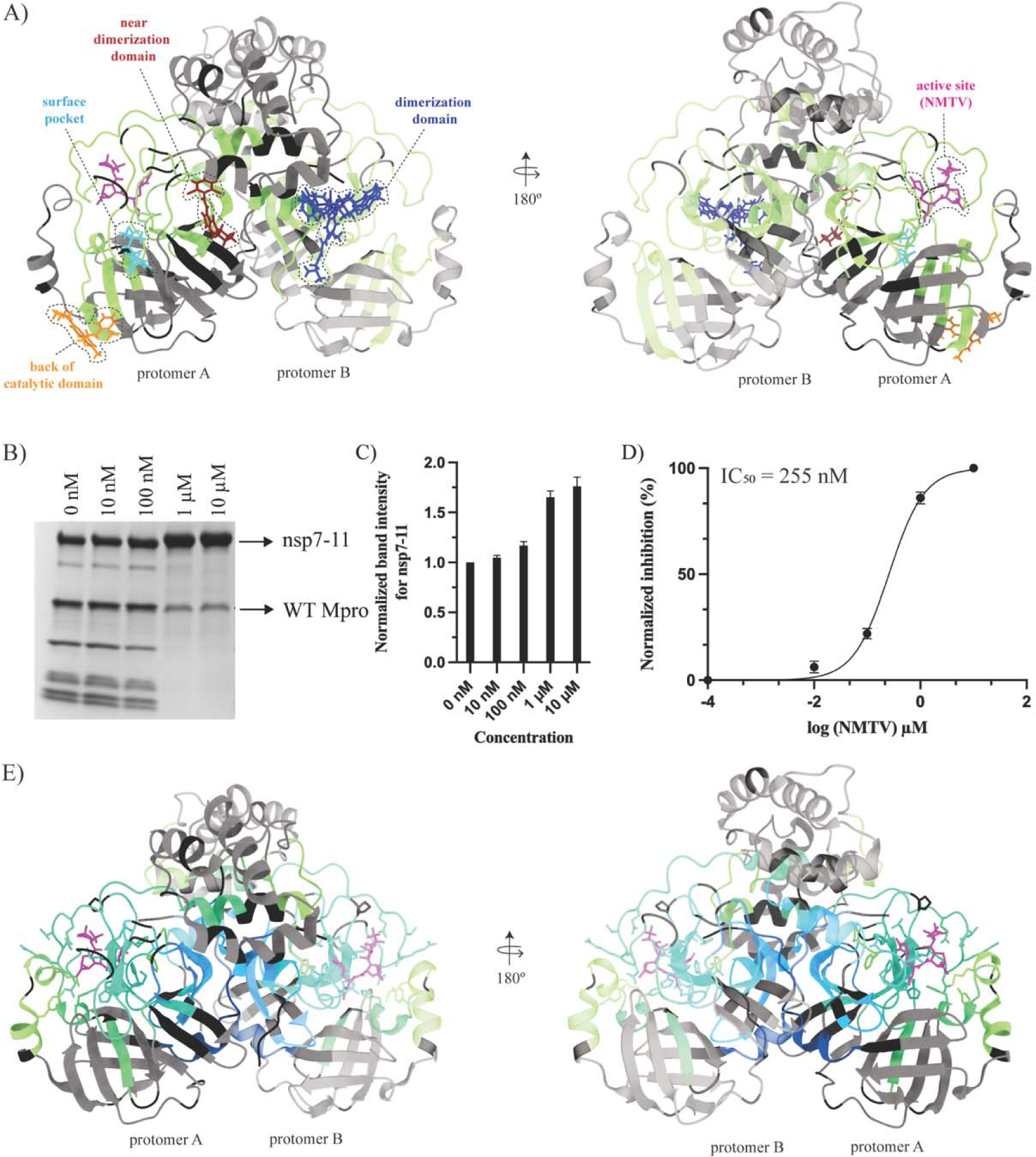
Inhibition of polyprotein processing by small molecules. (A) Mapping of small molecules or fragments onto the structure of Mpro (PDB 7DVY) that is colored according to the consolidated differential percent deuterium uptake values shown in **Fig. 2D**. Binders are shown as stick models and color-coded by the region of Mpro they interact with. See **Table S4** for more information regarding binders. (B) SDS-PAGE gel showing proteolytic processing of nsp7-11 by Mpro in the presence of increasing nirmatrelvir (NMTV) concentrations (0-10 µM) for 24 h. (C) Inhibition of NMTV is shown by plotting the normalized band intensities of the nsp7-11 substrate vs. NMTV concentrations. (D) Dose-response curve of NMTV inhibition of Mpro. The IC_50_ value was calculated from three independent replicates. (E) Differential HDX-MS results for Mpro in the presence and absence of NMTV overlaid onto the structure of Mpro with NMTV (PDB 7RFW).

NMTV, as expected, showed comparable inhibition with an IC_50_ value of 255 nM with the full polyprotein substrate *in vitro* **(Fig. 6B-D)** (*29*). None of the non-active site binders displayed significant inhibition of the enzymatic activity of Mpro **(Fig. S15A)**. On the contrary, climbazole and pelitinib showed activation of Mpro activity in our assay conditions, despite the last presenting an EC_50_ = 1.25 μM and moderate cytotoxicity (*32*).

Next, we analyzed by differential HDX-MS the effect of ligand binding on Mpro. Interestingly, NMTV was the only compound that showed significant change in Mpro solvent exchange behavior **(Fig. 6E, S15B, Table S1)**. The lack of observed change in solvent exchange may be due to experimental limitations on studying interactions of weak binders by HDX-MS (*53, 54*). The NMTV interaction footprint on Mpro demonstrates strong protection from solvent exchange in the active site, in agreement with mechanism of action of NMTV forming a reversible covalent thioimidate adduct with the catalytic C145 (*29*). These results closely resemble the nsp7-10/11 interaction footprint **(Fig. 2 and 6A)**, as we observed protection in the active site of Mpro upon interaction with the polyprotein. Additionally, the nsp7-10/11 footprint showed protection from solvent exchange in residues V77-L89 not found in the presence of NMTV. These residues are located on the back of the catalytic domain, near residues K61 and S62 which form inter-Mpro-nsp7-11 crosslinks and are thus likely stemming from more transient interaction of the Mpro with the polyprotein away from the active site.

## Discussion

In this work, we have studied the processing of CoV-2 nsp7-10/nsp7-11 polyproteins by Mpro. As expected by the high degree of amino acid conservation, we have seen that CoV-2 polyprotein processing is almost, if not, identical to that observed for CoV-1 (*22*). The cleavage order deduced from the gel analysis is also supported by results from the pulsed HDX-MS experiment. The destabilization observed in the pulsed HDX-MS in the nsp9 C-terminal region at the first time point (30 min) suggests cleavage and release of nsp7-9 to increase solvent exposure of the nsp9 C-terminal region. This is consistent with the fact that SDS-PAGE gel shows an intermediate nsp7-9 polyprotein observed at 30 min-1 h suggesting that the nsp9-10 junction is the first cleavage site. As shown in the literature for SARS-CoV (*22*), the order of processing cannot be directly inferred from the substrate specificity of Mpro with peptides mimicking the cleavage junctions as the conformation and accessibility of the substrate polyprotein(s) is critical to regulating the process.

Whether the same *in vitro* order of cleavages occur during viral replication is unknown. However, several lines of evidence support this concept. Several studies have detected the nsp4- nsp10/11 polyprotein intermediate in MHV-infected cells (*13, 16, 55*). Recently, in CoV-2 infected cells, the identification of viral cleavage sites at nsp4, nsp8-9, and nsp10-12 junctions at different post-infection time points is also consistent with such a polyprotein intermediate (3). Reverse genetics studies with MHV infected cells (8, 16, 55) also provide support for their essential role in the viral replication cycle. As shown in MHV, the processing order of the nsp7- 10 region is crucial for viral replication: either domain deletions or switching, and cleavage site mutations were lethal to the virus replication, with the exception being inactivation of the nsp9-10 cleavage site, which yielded an attenuated mutant virus (8).

Additionally, the nsp7-11 and nsp7-8 processing results indicate the presence of the nsp7- 8 intermediate even after 24 hours of exposure to Mpro. It is not known whether this longer-lived intermediate could have some functional or essential role in the viral cycle; further suppression of nsp7-8 maturation could represent a unique drug target. The existence of potent maturation inhibitors in HIV has validated this concept as a plausible strategy; bevirimat, the lead for this class, binds to the CA/SP1 junction of the Gag polyprotein and hinders its cleavage: this junction (similarly to the nsp7-8 junction) is in a dynamic helix-to-coil equilibrium and binding of bevirimat stabilizes the helical conformation (*56–58*). Regardless, it should be noted that, as labeling techniques used for microscopy cannot distinguish between mature nsps and polyprotein intermediates, chemical probes specifically targeting the nsp7-8 junction could help in further elucidation of the role of polyproteins during the CoV cycle.

The critical observation that the studied polyproteins have similar deuterium incorporation profiles as the individual proteins (and thus share similar structural elements) led us to conclude that the individual nsps do not undergo large structural rearrangements following cleavage by Mpro. This permitted the use of an integrative structural biology approach combining modeling and experimental methodologies to elucidate the structural basis for the order of CoV-2 polyprotein processing. The structural predictions of nsp7-11 polyprotein using the I-TASSER software provided us with an ensemble of 10 models with four representative conformations. Overall, none of the four groups satisfy all the experimental HDX-MS, XL-MS, and SAXS data, suggesting that the nsp7-11 polyprotein is highly dynamic and samples multiple conformations. While SAXS and HDX-MS capture the extended nsp8 helix conformation represented by Group A, XL-MS data are more consistent with the more globular protein conformations seen in Group B and Group C. The surface-accessible areas of the cleavage junctions and secondary structure element analysis of the nsp7-11 polyprotein suggested that Groups B and C (comprising six out of the ten models of the ensemble) might represent the polyprotein conformations in better agreement with the processing order we determined experimentally (*e.g.*, more accessible and disordered nsp9-10 junction in comparison with a more structured and hindered nsp7-8 junction). On the other hand, the four models comprising Groups A and D showcase the conformational adaptability of the polyprotein: in these models, the nsp7-8 junction is more exposed and unstructured, thus accessible for cleavage. Overall, the nsp7-11 model ensemble recapitulates the need for viral polyproteins to adopt different conformations during the replication cycle, i.e., metamorphic proteins (*59, 60*), given the strict genetic economy of RNA viruses.

The HDX-MS footprint and XL-MS of the Mpro:nsp7-11 complex reveal the importance of the “incognito” part of the polyprotein—the part of the polyprotein excluding the junctions captured in Mpro:substrate peptidic structures (*33–35*)—in processing. While we see that binding to Mpro substantially stabilizes the nsp8 portion of nsp7-11 **(Fig. 2A-B and 2E)**, positioning of the polyprotein may be such that either the polyprotein binds to the active site of one Mpro protomer and wraps around to make contact with the back side of the catalytic domain of that same protomer, or the polyprotein binds to the active site of one protomer and sits on top of the back side of the catalytic domain of the other protomer **(Fig. 7)**. The HDX-MS footprint on C145A Mpro shows that the binding to the active site is strongest, while the binding outside of it is more transient **(Fig. 2C** and **2D**, respectively). This could be because the polyprotein can adopt four different conformations that can interact differently with the catalytic domain and depending on which junction is interacting with the active site **(Fig. 2A-B),** highlighting the transitoriness of these interactions. The lack of Mpro inhibition by the non-active site surface binders also hints to this more transient nature **(Fig. S15)**. Nevertheless, despite this more transient nature, these interactions may be important in setting the conformation of the junctions for cleavage, as aforementioned.

**Fig 7.**
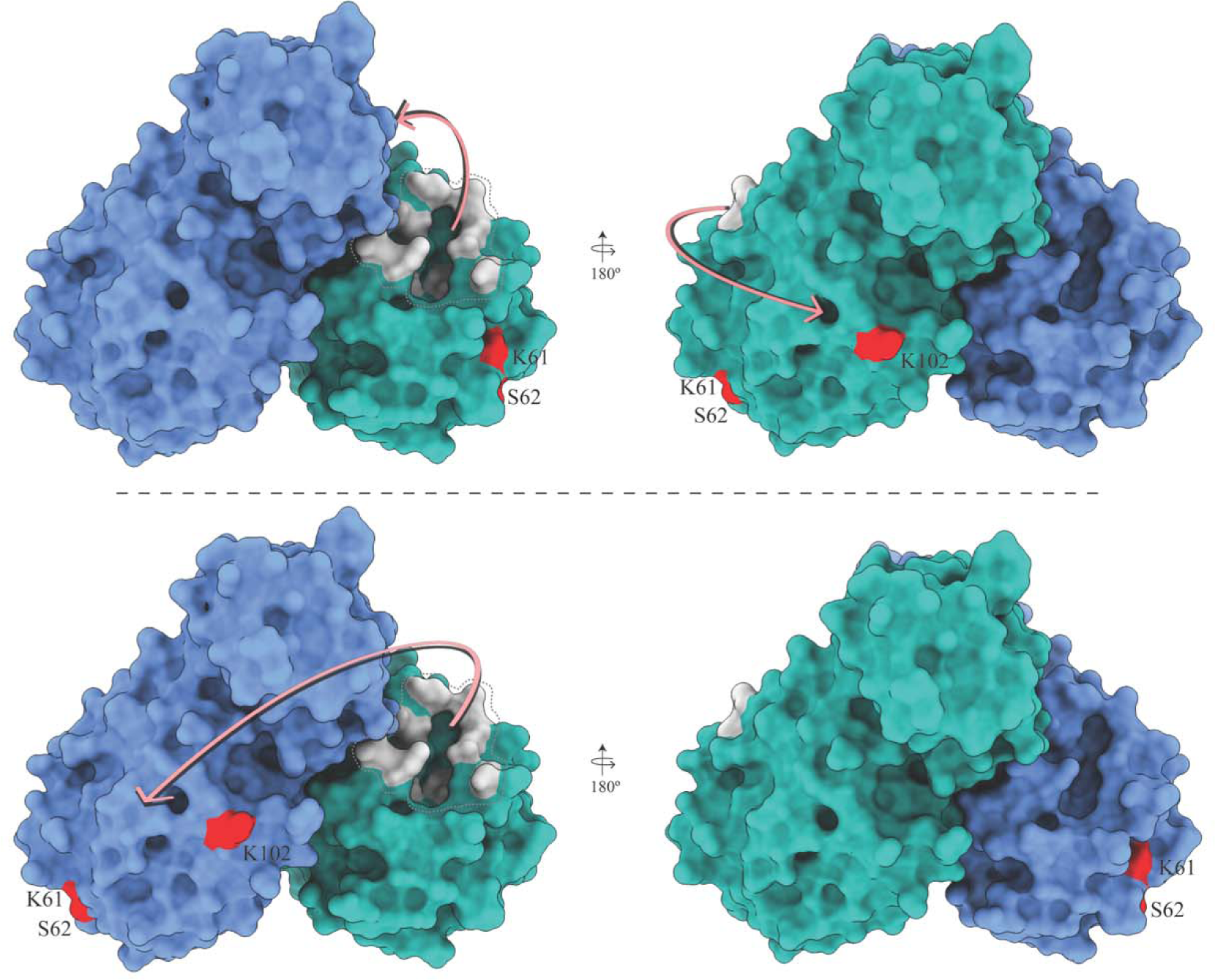
Schematic representation of the nsp7-11 polyprotein substrate binding to Mpro. Polyprotein binding to the active site of one Mpro protomer and wrapping around to contact the back side of the catalytic domain of that same protomer *(upper panel).* Polyprotein binding to the active site of one protomer and sitting on top of the back side of the catalytic domain of the other protomer *(lower panel).* The two protomers of Mpro are shown in blue and cyan. Protein residues shown in gray are the active site residues and residues in red are sites of inter-protein crosslinking with the polyprotein.

The HDX-MS footprint of Mpro:nsp7-11 also reveals significant protection in the Mpro dimerization interface area, especially near the Mpro C-terminus **(Fig. 2D and 6**) and suggesting that nsp7-11 binding stabilizes the Mpro dimer. In this sense, El-Baba and colleagues (*61*) identified that fragment JGY—discovered through crystallographic fragment screening and binding in the dimer interface (*30*)—destabilized the Mpro dimer and showed ∼35% inhibition of the rate of processing at 100 μM. Along the same lines, Sun and colleagues (*62*) discovered a nanobody, NB2B4, which binds the C-terminal domain of monomeric Mpro (PDB 7VFB) and inhibits activity with an IC_50_ ∼150 nM. Thus, destabilization of the Mpro catalytic dimer may contribute to the mechanism of inhibition. Combined with the lack of Mpro inhibition by the non- active site surface binders **(Fig. S15)**, allosteric inhibition of Mpro may only be efficiently achieved by interface binders destabilizing the Mpro dimer. Binding in areas on the surface may not distort the active site of Mpro, which is by nature very malleable (accommodating 11 different junctions *in virio*) (*25, 34*).

While the impressive crystallographic small molecule repurposing campaign (*32*) has provided valuable hits and probes along with antiviral activity testing, enzymatic inhibition was not reported by this study. As reviewed for remdesivir (*63*), the value of mechanistic, and enzymatic inhibition studies (alongside antiviral studies) is paramount, because it provides a logical path for developing direct-acting antivirals. The best example in the current case is pelitinib, which given its strong antiviral activity was portrayed as an allosteric inhibitor. Our studies show that it might be an allosteric activator. We hypothesize that this activation might be due to the stabilization of the Mpro dimer **(Fig. S15A)**. To understand whether its antiviral activity is due to an off-target effect [pelitinib has low inhibition of PLpro in enzymatic inhibition assays, (*64*)] or due to dysregulation of viral maturation [as seen for efavirenz acceleration of Gag-Pol processing in HIV (*65*)], more experiments are required.

In summary, this study describes the structural basis of the order of Mpro processing of the essential nsp7-10/11 segment, the importance of the more transient interactions of the substrate to Mpro for proper positioning and catalysis and provides a mechanistic validation of allosteric inhibition. In conclusion, our results give structural insights into CoV-2 polyproteins which will help us in understanding the structure-function relationships, drug design, and the fundamental biology of polyprotein activity and processing in CoV-2.

## Materials and Methods

### Reagents and Plasmids

Unless otherwise specified, all chemicals and reagents were purchased from Sigma- Aldrich (St. Louis, MO). Formic acid, trifluoroacetic acid, and UHPLC-grade solvents were purchased from ThermoFisher. The active site inhibitor NMTV and non-active site binder RS102895 were purchased from MedChemExpress. AT7519 and Climbazole were purchased from Selleck Chemicals. PD168568 was purchased from Tocris Bioscience. Pelitinib was purchased from BioVision. The pGEX-6P-1-nsp5 (or Mpro) plasmid was a kind gift from Dr. Martin Walsh, Diamond Light Source. pGBWm4046979 (coding for full-length nsp7, NCBI Reference Sequence: YP_009725303.1, codon-optimized, with an initial Met and a cleavable C- terminal TEV 6x-His tag was a gift from Ginkgo Bioworks (Addgene plasmid 145611; http://n2t.net/ addgene:145611; RRID: Addgene_145611). pGBWm4046852 (coding for full- length nsp8, NCBI Reference Sequence: YP_009725304.1, codon-optimized, with an initial Met and a cleavable C-terminal TEV 6x-His tag) was a gift from Ginkgo Bioworks (Addgene plasmid 145584; http://n2t.net/ addgene:145584; RRID: Addgene_145584). The pET-28a-nsp9 gene was obtained from BEI Resources (NR-53501). The gene encoding SARS-CoV-2 nsp10 was cloned into the pGEX-6P-1 vector to generate an expression construct containing an N-terminal GST tag and an HRV 3C protease cleavage site (GST_3C_Nsp10). Plasmids for codon-optimized pET-28a- His_6_-nsp7-8 and pET-28a-His_6_-nsp7-11 (with an HRV 3C protease cleavage site between the 6x- His tag and the coding sequence) were obtained from GenScript (Piscataway, NJ). Primers used for cloning and mutagenesis, as well as plasmid sequences, are available upon request. HRV 3C and TEV proteases were recombinantly expressed using in-house plasmids.

### Protein Expression and Purification

WT Mpro was produced with native N- and C-termini, as described in (*66*). The pGEX- 6P-1-nsp5 expression plasmid was transformed into *E. coli* Rosetta gami competent cells and cultured in LB media at 37 °C with 100 μg/mL ampicillin. Next day, the culture was diluted 1:100 into 1 L LB media supplemented with 100 μg/mL ampicillin. The cells were grown to an OD_600_ (optical density at 600 nm) = 0.8 before being induced with 1 mM IPTG at 16 °C. After 10 h of induction, the cells were collected by centrifugation at 7,200 x *g* for 10 min and stored in -80 °C.

The cell pellet was resuspended in 50 mM Tris pH 8.0, 300 mM NaCl, 5 mM imidazole, and 1 mM TCEP followed by sonication and centrifugation at 30,000 x *g* for 60 min. The cleared lysate was loaded on a Ni-NTA affinity column (Qiagen). The bound proteins were first washed with lysis buffer and then with the lysis buffer supplemented with 20 mM imidazole to remove non- specific proteins. Mpro was eluted with 300 mM imidazole in the lysis buffer and then purified by size-exclusion chromatography using a pre-packed Superose 6 Increase 10/300 GL column (GE Healthcare Life Sciences) equilibrated in 50 mM Tris pH 8.0, 300 mM NaCl, and 1 mM TCEP. The fractions containing the pure protein were pooled, concentrated, and stored in –80 °C.

nsp7 and nsp8 were produced as described in our earlier work (*37*). nsp9 was purified using the protocol described in (*67*). Overall, the plasmid was first transformed into *E. coli* BL21- CodonPlus (DE3)-RIL cells and then grown in LB media with 50 μg/mL kanamycin at 37 °C. The cells were grown to an OD_600nm_ 1.0 before being induced with 0.5 mM IPTG. After 4 h of induction, the cells were collected by centrifugation at 7,200 x *g*. The cells were resuspended in a lysis buffer (20 mM HEPES pH 7.0, 150 mM NaCl, 20 mM imidazole, 2 mM MgCl_2_, and 0.5 mM TCEP). The cells were lysed by sonication in the presence of 1 mg of lysozyme and then centrifuged at 10,000 x *g* for 20 min. The cleared lysate was loaded on a Ni-NTA affinity column (Qiagen) and the column was then washed with the lysis buffer and 50 mM imidazole buffer. The His-tagged protein was eluted with 400 mM imidazole in the lysis buffer. The His-tag was cleaved by incubating the protein with HRV 3C protease overnight at 4 °C. After digestion, the protein was passed through a second Ni-NTA column to remove the 3C protease and the residual un-cleaved protein. The protein sample was then purified by size-exclusion chromatography using a pre-packed Superose 6 Increase 10/300 GL (GE Healthcare Life Sciences) equilibrated in 20 mM HEPES pH 7.0, 150 mM NaCl, 2 mM MgCl_2,_ and 0.5 mM TCEP. Pure protein-containing fractions were concentrated and stored at -80 °C after snap freezing.

A single colony of *E. coli* BL-CodonPlus (DE3)-RIL (Agilent Technologies) carrying the GST_3C_-nsp10 was used to inoculate 50 mL of LB media containing the appropriate antibiotics (100 μg/mL carbenicillin and 25 μg/mL chloramphenicol). This seeding culture was grown overnight in a shaking incubator at 37 °C. The seeding cultures were then used to inoculate 1 L expression cultures containing the appropriate antibiotics to an initial OD_600_ of 0.2 and grown in a shaking incubator at 37 °C to an OD_600_ of 0.6. The temperature was reduced to 16 °C, and protein expression was induced at an OD_600_ of 0.9 with the addition of 0.1 mM IPTG. The expression cultures were harvested after 16 hours by centrifugation for 30 min at 2555 x *g*, followed by flash-freezing and storage at -80 °C. All centrifugation steps were performed at 4 °C. The cell pellet from 1 L of expression culture was resuspended in lysis buffer (50 mM Tris, 300 mM NaCl, 5 mM β-mercaptoethanol, 4 mM MgSO_4_, 10 % volume/volume glycerol, pH 8.0), at a ratio of 5 mL lysis buffer to 1 g cell paste and thawed on ice. The cells were lysed by sonication on ice for 8 minutes, and the cellular debris was separated from the soluble lysate by centrifugation for 30 min at 48,000 x *g*. The volume of the soluble lysate was measured, and an equal volume of saturated ammonium sulfate was added to achieve 50% saturation, followed by overnight incubation at 4 °C. The soluble fraction was separated by centrifugation for 30 min at 24,000 x *g* and discarded. The pellet was resuspended in 10 mL of lysis buffer, and 100 μL of polyethyleneimine (5% w/v) was added in a dropwise fashion. The insoluble material was removed by centrifugation for 30 minutes at 24,000 x *g*. The supernatant was decanted and added to 2 mL of Glutathione Sepharose 4 FF (Cytiva) affinity medium which had been pre-equilibrated with lysis buffer. Batch binding was performed on an orbital rotator at 4 °C for 4 h, and the unbound protein was removed using gravity-flow chromatography and washed with 20 mL of lysis buffer. GST_3C_-nsp10 was cleaved on-column with the addition of 8 mL of lysis buffer containing 0.2 mg/mL HRV 3C protease, and incubation on an orbital rotator at 4 °C overnight. The cleaved nsp10 was collected in the flow-through and wash fractions, concentrated to 198 μM using an Amicon Ultra centrifugal filter (Millipore Sigma), and stored at -80 °C.

The nsp7-8 and nsp7-11 polyprotein genes were transformed into *E. coli* BL21-CodonPlus (DE3)-RIL cells and grown overnight on an LB-agar plate containing 50 μg/mL kanamycin. A single colony was picked from the plate and inoculated into LB media with 50 μg/mL kanamycin. The culture was grown overnight at 37°C. Next morning, the starter culture was diluted 1:500 into the LB media. The cells were grown at 37 °C until OD_600_ ∼ 1.6 was reached. The culture was then allowed to cool for an hour at 20 °C with continuous shaking after which it was induced with 1 mM IPTG. After overnight incubation, the cells were collected by centrifugation at 7,200 x *g*. The cell pellet was resuspended in lysis buffer (50 mM Tris pH 8.0, 500 mM NaCl, 20 mM imidazole, 5% glycerol, 10 mM CHAPS, 1 mM TCEP) supplemented with 1 µM leupeptin, 1 µM pepstatin, and 1 mM PMSF. The cell suspension was lysed by sonication and clarified by centrifugation at 30,000 x *g* at 4 °C for an hour. The supernatant was loaded on a Ni-NTA affinity column (Qiagen), pre-equilibrated with the lysis buffer. The column was first washed with lysis buffer and then with 50 mM imidazole in lysis buffer. Homemade HRV 3C protease in the buffer containing 50 mM Tris pH 8.0, 500 mM NaCl, 20 mM imidazole, 5% glycerol, 1 mM TCEP, was added to perform on-column cleavage of the 6x-His tag at 4°C. The digested protein was eluted from the column and passed through a second Ni-NTA column. This reverse Ni-NTA step is performed to remove residual 3C protease and uncleaved protein. The protein was further purified by ion-exchange chromatography using the HiTrap Heparin HP column (GE Healthcare Life Sciences) and a Mono Q anion exchange column (16/10; GE Healthcare Life Sciences) using gradient elution from 150 mM to 2 M NaCl. The protein sample was then purified by size-exclusion chromatography using a pre-packed Superose 6 Increase 10/300 GL (GE Healthcare Life Sciences) equilibrated in 50 mM Tris pH 8.0, 500 mM NaCl, 5% glycerol, and 1 mM TCEP. Pure protein-containing fractions were pooled together, concentrated, and stored at -80 °C.

### Proteolysis assays with nsp7-8 and nsp7-11 polyprotein substrates

WT Mpro was used to carry out cleavage assays with the nsp7-8 and nsp7-11 polyprotein substrates. The *in vitro* cleavage reaction was performed by incubating the polyproteins with Mpro WT (nsp7-11:Mpro molar ratio was 6:0.5, in µM; nsp7-8:Mpro molar ratio was 5:0.5, in µM) at room temperature in the assay buffer: 50 mM Tris pH 7.5, 150 mM NaCl, and 1mM DTT. The reaction was stopped at various time points by the addition of 4X stop buffer (277.8 mM Tris-HCl pH 6.8, 44.4% glycerol, 4.4% SDS, 0.02% bromophenol blue). The samples were then denatured at 95 °C for 5 min and assessed on a gradient SDS-PAGE gel. The bands for the full-length substrates, intermediate products, and the final cleavage products were cut and confirmed by mass spectrometry. In-gel trypsin digestion was performed on the gel bands and LC-MS/MS was carried out on them (see SM for MS experimental details).

### In vitro assessment of the effects of small molecules on Mpro activity

The stock solutions of all the Mpro binders were made in DMSO. They were diluted in the assay buffer and pre-incubated for 30 min at room temperature with Mpro WT before starting the reaction. The nsp7-11 polyprotein substrate was then added to the reaction at 6 µM. The reaction was stopped after 24 h. After denaturing, the samples were then run on the SDS-PAGE gel. The effect of small molecules on Mpro activity was assessed by observing the amount of substrate (nsp7-11) present after 24 h. The gel band intensity for nsp7-11 was calculated using ImageJ software (https://imagej.nih.gov/ij/index.html) and plotted against the concentration of binders using the GraphPad Prism Version 9.3.1 (GraphPad Software, La Jolla California USA, www.graphpad.com). The IC_50_ value calculation for NMTV was also done using the GraphPad Prism Version 9.3.1.

### Crosslinking mass spectrometry (XL-MS)

#### Sample preparation

For DSSO (disuccinimidyl sulfoxide) (ThermoFisher) crosslinking reactions, individual protein and protein-protein complexes were diluted to 10 µM in crosslinking buffer (50 mM HEPES pH 8.0, 500 mM NaCl, 1 mM TCEP) and incubated for 30 min at room temperature prior to initiating the crosslinking reaction. DSSO crosslinker was freshly dissolved in crosslinking buffer to a final concentration of 75 mM before being added to the protein solution at a final concentration of 1.5 mM. The reaction was incubated at 25 °C for 45 or 90 min, then quenched by adding 1 µL of 1.0 M Tris pH 8.0 and incubating an additional 10 min at 25°C. Control reactions were performed in parallel without adding the DSSO crosslinker. All crosslinking reactions were carried out in three replicates. The presence of crosslinked proteins was confirmed by comparing to the no-crosslink negative control samples using SDS-PAGE and Coomassie staining. The remaining crosslinked and non-crosslinked samples were separately pooled and then precipitated using methanol and chloroform. Dried protein pellets were resuspended in 12.5 µL of resuspension buffer (50 mM ammonium bicarbonate, 8 M urea, pH 8.0). ProteaseMAX (Promega - V5111) was added to 0.02% and the solutions were mixed on an orbital shaker operating at 400 rpm for 5 min. After resuspension, 87.5 µL of digestion buffer (50 mM ammonium bicarbonate pH 8.0) was added. Protein samples were reduced by adding 1 µL of 500 mM DTT followed by incubation of the protein solutions on an orbital shaker operating at 400 rpm at 56 °C for 20 minutes. After reduction, 2.7 µL of 550 mM iodoacetamide was added and the solutions were incubated at room temperature in the dark for 15 min. Reduced and alkylated protein solutions were digested overnight using trypsin at a ratio of 1:150 (w/w trypsin:protein) at 37°C. Peptides were acidified with 1% trifluoroacetic acid (TFA) and then desalted using C18 ZipTip® (Millipore cat # ZTC18 5096). Dried peptides were resuspended in 10 µL of 0.1% TFA in water. Samples were then frozen and stored at -20 °C until LC-MS analysis.

### Liquid Chromatography and Mass Spectrometry

500 ng of sample was injected (triplicate injections for crosslinked samples and duplicate injections for control samples) onto an UltiMate 3000 UHP liquid chromatography system (Dionex, ThermoFisher). Peptides were trapped using a μPAC C18 trapping column (PharmaFluidics) using a load pump operating at 20 µL/min. Peptides were separated on a 200 cm μPAC C18 column (PharmaFluidics) using a linear gradient (1% Solvent B for 4 min, 1-30% Solvent B from 4-70 min, 30-55% Solvent B from 70-90 min, 55-97% Solvent B from 90-112 min, and isocratic at 97% Solvent B from 112-120 min) at a flow rate of 800 nL/min. Gradient Solvent A contained 0.1% formic acid and Solvent B contained 80% acetonitrile and 0.1% formic acid. Liquid chromatography eluate was interfaced to an Orbitrap Fusion Lumos Tribrid mass spectrometer (ThermoFisher) with a Nanospray Flex ion source (ThermoFisher). The source voltage was set to 2.5 kV, and the S-Lens RF level was set to 30%. Crosslinks were identified using a previously described MS2-MS3 method (*68*) with slight modifications. Full scans were recorded from m/z 150 to 1500 at a resolution of 60,000 in the Orbitrap mass analyzer. The AGC target value was set to 4e5, and the maximum injection time was set to 50 ms in the Orbitrap. MS2 scans were recorded at a resolution of 30,000 in the Orbitrap mass analyzer. Only precursors with charge state between 4-8 were selected for MS2 scans. The AGC target was set to 5e4, a maximum injection time of 150 ms, and an isolation width of 1.6 m/z. CID fragmentation energy was set to 25%. The two most abundant reporter doublets from the MS2 scans with a charge state of 2-6, a 31.9721 Da mass difference, and a mass tolerance of ±10 ppm were selected for MS3. The MS3 scans were recorded in the ion trap in rapid mode using HCD fragmentation with 35% collision energy. The AGC target was set to 2e4, the maximum injection time was set for 200 ms, and the isolation width set to 2.0 m/z.

### Data Analysis

To identify crosslinked peptides, Thermo.Raw files were imported into Proteome Discoverer 2.5 (ThermoFisher) and analyzed via XlinkX algorithm (*69*) using the MS2_MS3 workflow with the following parameters: MS1 mass tolerance—10 ppm; MS2 mass tolerance— 20 ppm; MS3 mass tolerance—0.5 Da; digestion—trypsin with four missed cleavages allowed; minimum peptide length of 4 amino acids, fixed modification—carbamidomethylation (C); variable modification—oxidation (M); and DSSO (K, S, T, Y). The XlinkX/PD Validator node was used for crosslinked peptide validation with a 1% false discovery rate (FDR). Identified crosslinks were further validated and quantified using Skyline (version 19.1) (*70*) using a previously described protocol (*71*). Crosslink spectral matches found in Proteome Discoverer were exported and converted to sequence spectrum list format using Excel (Microsoft). Crosslink peak areas were assessed using the MS1 full-scan filtering protocol for peaks within 8 min of the crosslink spectral match identification. Peak areas were assigned to the specified crosslinked peptide identification if the mass error was within 10 ppm of the theoretical mass, the isotope dot product was greater than 0.95, and if the peak was not found in the non-crosslinked negative control samples. The isotope dot product compares the distribution of the measured MS1 signals against the theoretical isotope abundance distribution calculated based on the peptide sequence. Its value ranges between 0 and 1, where 1 indicates a perfect match (*72*). Pair-wise comparisons were made using the ‘MSstats’ package (*73*) implemented in Skyline to calculate relative fold changes and significance. Significant change thresholds were defined as a log2 fold change less than -2 or greater than 2 and -log10 p-value greater than 1.3 (p-value less than 0.05). Visualization of proteins and crosslinks was generated using xiNET (*74*).

The data have been deposited to the ProteomeXchange Consortium via the PRIDE (*75*) partner repository with the dataset identifier PXD033748.

**Project Name:** Biochemical and structural insights into SARS-CoV-2 polyprotein processing by Mpro (XL-MS)

**Project accession:** PXD033748 Reviewer account details:

**Password:** oKi04fUZ

### Hydrogen-deuterium exchange mass spectrometry (HDX-MS)

#### Peptide identification

Peptides were identified using tandem MS (MS/MS) experiments performed on a QExactive (Thermo Fisher Scientific, San Jose, CA) over a 70 min gradient. Product ion spectra were acquired in a data-dependent mode and the five most abundant ions were selected for the product ion analysis per scan event. The MS/MS *.raw data files were converted to *.mgf files and then submitted to MASCOT (version 2.3 Matrix Science, London, UK) for peptide identification. The maximum number of missed cleavages was set at 4 with the mass tolerance for precursor ions +/- 0.6 Da and for fragment ions +/- 8 ppm. Oxidation to methionine was selected for variable modification. Pepsin was used for digestion and no specific enzyme was selected in MASCOT during the search. Peptides included in the peptide set used for HDX detection had a MASCOT score of 20 or greater. The MS/MS MASCOT search was also performed against a decoy (reverse) sequence and false positives were ruled out if they did not pass a 1% false discovery rate.

### Pulse labeling

The nsp7-10 polyprotein at 10 μM concentration was incubated with WT Mpro at 1:1 molar ratio and 5 μL aliquots of the cleavage reaction were removed at 600, 1800, 3600, 14400, and 86400 s. Aliquots were mixed with 20 μL of deuterated (D_2_O-containing) buffer (50 mM HEPES, 500 mM NaCl, 1 mM TCEP, pD 8.4) and incubated on ice for 30 s. Deuterated samples were quenched with 25 μL quench solution (5 M urea, 1% TFA, pH 2) and immediately flash frozen and stored until ready for direct inject MS analysis.

### Continuous labeling

Experiments with continuous labeling were carried out on a fully automated system (CTC HTS PAL, LEAP Technologies, Carrboro, NC; housed inside a 4 °C cabinet) as previously described (*76*) with the following modifications. For differential HDX, protein-protein complexes were pre-formed and allowed to incubate 30 min at room temperature prior to analysis. The reactions (5 μL) were mixed with 20 μL of deuterated (D_2_O-containing) buffer (50 mM HEPES, 500 mM NaCl, 1 mM TCEP, pD 8.4) and incubated at 4 °C for 0 s, 10 s, 30 s, 60 s, 900 s, or 3600 s. Following on-exchange, unwanted forward- or back-exchange was minimized, and the protein was denatured by the addition of 25 μL of a quench solution (5 M urea, 1% TFA, pH 2.0) before being immediately passed along for online digestion.

### HDX-MS analysis

Samples were digested through an immobilized pepsin column (prepared in-house) at 50 μL/min (0.1% v/v TFA, 4 °C) and the resulting peptides were trapped and desalted on a 2 mmL×L10 mm C8 trap column (Hypersil Gold, ThermoFisher). The bound peptides were then gradient-eluted (4-40% CH_3_CN v/v and 0.3% v/v formic acid) on a 2.1 mm × 50 mm C18 separation column (Hypersil Gold, ThermoFisher) for 5 min. Sample handling and peptide separation were conducted at 4 °C. The eluted peptides were then subjected to electrospray ionization directly coupled to a high-resolution Orbitrap mass spectrometer (QExactive, ThermoFisher).

### Data rendering

The intensity weighted mean m/z centroid value of each peptide envelope was calculated and subsequently converted into a percentage of deuterium incorporation. This is accomplished by determining the observed averages of the undeuterated and fully deuterated spectra using the conventional formula described elsewhere (*77*). The fully deuterated control, 100% deuterium incorporation, was calculated theoretically, and corrections for back-exchange were made on the basis of an estimated 70% deuterium recovery and accounting for 80% final deuterium concentration in the sample (1:5 dilution in deuterated buffer). Statistical significance for the differential HDX data is determined by an unpaired t-test for each time point, a procedure that is integrated into the HDX Workbench software (*78*).

The HDX data from all overlapping peptides were consolidated to individual amino acid values using a residue averaging approach. Briefly, for each residue, the deuterium incorporation values and peptide lengths from all overlapping peptides were assembled. A weighting function was applied in which shorter peptides were weighted more heavily and longer peptides were weighted less. Each of the weighted deuterium incorporation values were then averaged incorporating this weighting function to produce a single value for each amino acid. The initial two residues of each peptide, as well as prolines, were omitted from the calculations. This approach is similar to that previously described (*79*).

Deuterium uptake for each peptide is calculated as the average of %D for all on-exchange time points and the difference in average %D values between the unbound and bound samples is presented as a heat map with a color code given at the bottom of the figure (warm colors for deprotection and cool colors for protection). Peptides are colored by the software automatically to display significant differences, determined either by a >5% difference (less or more protection) in average deuterium uptake between the two states, or by using the results of unpaired t-tests at each time point (p-value < 0.05 for any two time points or a p-value < 0.01 for any single time point). Peptides with non-significant changes between the two states are colored gray. The exchange at the first two residues for any given peptide is not colored. Each peptide bar in the heat map view displays the average Δ %D values, associated standard deviation, and the charge state. Additionally, overlapping peptides with a similar protection trend covering the same region are used to rule out data ambiguity.

The data have been deposited to the ProteomeXchange Consortium via the PRIDE (*75*) partner repository with the dataset identifier PXD033702 for the pulse labeling HDX-MS experiment and PXD033698 for continuous labeling HDX-MS experiments.

**Project Name:** Biochemical and structural insights into SARS-CoV-2 polyprotein processing by Mpro (HDX-MS continuous labeling)

**Project accession:** PXD033698 Reviewer account details:

**Password:** QgiXipcs

**Project Name:** Biochemical and structural insights into SARS-CoV-2 polyprotein processing by Mpro (HDX-MS pulse labeling)

**Project accession:** PXD033702 Reviewer account details:

**Password:** wHh4zEPB

### SEC-MALS-SAXS

Purified and concentrated nsp7-8 (8 mg/mL) and nsp7-11 (4 mg/mL) were used for data collection. SAXS was performed at BioCAT (beamline 18ID at the Advanced Photon Source, Chicago) with in-line size exclusion chromatography (SEC) to separate sample from aggregates and other contaminants thus ensuring optimal sample quality and multiangle light scattering (MALS), dynamic light scattering (DLS) and refractive index measurement (RI) for additional biophysical characterization (SEC-MALS-SAXS). The samples were loaded on a Superdex 200 Increase 10/300 GL column (Cytiva) run by a 1260 Infinity II HPLC (Agilent Technologies) at 0.6 mL/min. The flow passed through (in order) the Agilent UV detector, a MALS detector and a DLS detector (DAWN Helios II, Wyatt Technologies), and an RI detector (Optilab T-rEX, Wyatt). The flow then went through the SAXS flow cell. The flow cell consists of a 1.0 mm ID quartz capillary with ∼20 μm walls. A coflowing buffer sheath is used to separate samples from the capillary walls, helping prevent radiation damage (*80*). Scattering intensity was recorded using a Pilatus3 X 1M (Dectris) detector which was placed 3.69 m from the nsp7-11 sample giving us access to a q-range of 0.003 Å^-1^ to 0.35 Å^-1^ and 3.631 m from the nsp7-8 sample giving us access to a q-range of 0.0047 Å^-1^ to 0.35 Å^-1^. The data was reduced using BioXTAS RAW 2.0.3 (*81*). Buffer blanks were created by averaging regions flanking the elution peak and subtracted from exposures selected from the elution peak to create the I(q) vs q curves used for subsequent analyses. Molecular weights and hydrodynamic radii were calculated from the MALS and DLS data respectively using the ASTRA 7 software (Wyatt). Data analysis was carried out using the RAW software package for the determination of radius of gyration (Rg), P(r) distribution, particle maximum dimension (Dmax) parameters, and for qualitative flexibility analysis (through generation of Rg-Normalized Kratky and Guinier plots). Volumetric bead modeling was performed using the DAMMIN software package (*52*). The resulting bead models were averaged and filtered using the DAMAVER package (*82*), generating the final bead model reconstruction. The SAXS data are deposited in the SAXS database under the accession codes SASDPY2, SASDPZ2, SASDP23 and SASDP33.

### Structural integrative modeling using I-TASSER

For the structural predictions of the nsp7-11 polyprotein, an integrative modeling approach was employed. The I-TASSER server (*43*), which is an online source for automated protein structure prediction, was used to generate models of the polyproteins. A two-run approach was used to model the nsp7-11 polyprotein. Run 1 included the following inputs: i) amino acid sequence, ii) distance constraints from XL-MS, iii) nsp7-8 model as a template, and iv) secondary structure constraints for nsp7, nsp8, nsp9, and nsp10 as advised by HDX-MS to generate Models A1-2, D, and C1-C2 **(Fig. 3B-C).** Run 2 included: i) amino acid sequence, ii) distance constraints from XL-MS, iii) nsp7-8 as a template, and iv) secondary structure constraints for nsp8, nsp9, and nsp10 as advised by HDX-MS to generate Models A3 and B1-4 **(Fig. 3B-C).** Initial observation of the polyprotein by HDX-MS showed a similar pattern of deuterium uptake compared to the individual proteins **(Fig. 3A)**, suggesting that secondary structures within the polyprotein are likely to largely resemble the secondary structures of the mature nsps. Accordingly, this allowed us to delineate secondary structural constraints based on solved X-ray crystal structures of nsp7, nsp8, nsp9, and nsp10. We also used two nsp7-8 models we previously generated using a similar integrative modeling workflow to serve as additional structural templates since the HDX-MS footprint of the nsp7-8 polyprotein resembles the footprint of nsp7-8 in nsp7-11 **(Fig. 3A)**. The two nsp7-8 models were chosen based on their varying agreement with the XL-MS and HDX-MS data in order to limit bias from a particular experimental approach and sample the conformational landscape as thoroughly as possible **(see SM for nsp7-8 integrative modeling)**.

The ten nsp7-11 output models were assessed against the experimental (i) HDX-MS, (ii) XL-MS and the (iii) SAXS data: (i) Agreement of models to HDX-MS data was completed using HDXer which generated theoretical deuterium uptake values for the models to compare to experimental values (*49, 50*). Smaller RMSE indicates better agreement of models to experimental data. (ii) Crosslinks were mapped on the models using xiVIEW (doi: <u>10.1101/561829</u>) to calculate distances and determine percentage of crosslinks satisfied, *i.e.*, distances less than 30 Å. (iii) A theoretical scattering profile was generated for each model using the CRYSOL web interface (*83*). The theoretical scattering profile of each model was then fitted against the experimental scattering profile. Finally, the secondary structural elements and the solvent-accessible surface area of the junction sites for both polyproteins were also analyzed, and the results were compared with the limited proteolysis results to evaluate the physiological relevance of the structure in the context of polyprotein processing. The junction accessible area was calculated by the summation of the accessible area of four residues (P1, P2, P1’, P2’) at the junction site. The accessible surface area for each residue was calculated using VADAR (*84*). The integrative structures of nsp7-11 polyprotein have been deposited in the PDB-Dev databank under accession code PDBDEV_00000120. They are also provided in the SM as PyMOL sessions.

## Supporting information

SARS-CoV-2 nsp7-8 and nsp7-11 polyprotein integrative structures

## Acknowledgments

We thank Dr. Manju Narwal and Prof. Katsuhiko Murakami (Pennsylvania State University) for graciously providing protein samples for the C145A Mpro mutant and the nsp7-10 polyprotein and performing the proteolysis assay of nsp7-10 polyprotein with WT Mpro. We thank Bruce Pascal and Roberto Vera Alvarez (Omics Informatics LLC) for their help running the nsp7-11 and nsp7-8 models through the HDXer software. We thank Dr. Haiyan Zheng and Prof. Peter Lobel for performing the in-gel LC-MS/MS for the limited proteolysis assay of nsp7-11 polyprotein by Mpro. Molecular graphics and analyses were performed with: 1) UCSF Chimera, developed by the Resource for Biocomputing, Visualization, and Informatics at the University of California, San Francisco; and 2) PyMOL, (DeLano WL (2002) The PyMOL molecular graphics system. San Carlos, CA: DeLano Scientific).

## Author contributions

Each author’s contribution(s) to the paper are as follows:

Conceptualization: EA, PRG, FXR, RY, VVC, SKD

Methodology: RY, VVC, SKD, JJEKH, JT, JBH, FXR, RLS

Investigation: RY, VVC, SKD, RLS, SGS, EA, PRG, FXR

Visualization: RY, VVC, SKD, FXR

Supervision: EA, PRG, FXR

Writing—original draft: RY, VVC, FXR, EA, PRG

Writing—review & editing: all authors

## Data and materials availability

All data needed to evaluate the conclusions in the paper are present in the paper, including accession numbers to databases, and the Supplementary Materials. Correspondence and requests for materials should be addressed to Eddy Arnold, Patrick R. Griffin, and Francesc X. Ruiz.

## Supplementary Materials

Supplementary Text Figs. S1 to S15 Tables S1 to S4

## References

Other Supplementary Materials for this manuscript include the following:

PyMOL sessions with integrative structures of nsp7-8 and nsp7-11 polyproteins

