## Supplemental Material for "Biochemical and structural insights into SARS-CoV-2 polyprotein processing by Mpro"

**Affiliations**

**This PDF file includes:**

Supplementary Text

Figs. S1 to S15

Tables S1 and S4

References

**Other Supplementary Materials for this manuscript include the following:**

PyMOL sessions with integrative structures of nsp7-8 and nsp7-11 polyproteins

Supplementary Text

**Materials and Methods**

**Gel LC-MS/MS**

Samples (gel bands) were in-gel digested with sequencing grade trypsin (ThermoFisher) using standard protocol (1*,* 2) and analyzed by nano-LC-MS/MS.

***Nano-liquid chromatography-tandem mass spectrometry (LC-MS/MS)***

Samples were analyzed using a Q Exactive HF tandem mass spectrometer coupled to a Dionex Ultimate 3000 RLSCnano System (Thermo Scientific). Samples were loaded onto a fused silica trap column Acclaim PepMap 100, 75umx2cm (ThermoFisher). After washing for 5 min at 5 µl/min with 0.1% TFA, the trap column was brought in-line with an analytical column (Nanoease MZ peptide BEH C18, 130A, 1.7um, 75 um x 250 mm, Waters) for LC-MS/MS. Peptides were eluted using a segmented linear gradient from 4 to 90% B (A: 0.2% formic acid, B: 0.08% formic acid, 80% ACN): 4–15% B in 5 min, 15-50% B in 50 min, and 50-90% B in 15 min. Mass spectrometric data were acquired using a data-dependent acquisition procedure with a cyclic series of a full scan from 250-2000 with resolution of 120,000 followed by MS/MS (HCD, relative collision energy 27%) of the 20 most intense ions (charge +1 to +6) and a dynamic exclusion duration of 20 sec. Major peptides were manually confirmed.

***Database Search***

The peak list of the LC-MSMS were generated by Thermo Proteome Discoverer (v.2.1) into MASCOT Generic Format (MGF) and searched against custom supplied sequences plus a database composed of common lab contaminants using in house version of X!Tandem (3). Search parameters are as follows: fragment mass error: 20 ppm, parent mass error: +/- 7 ppm; fixed modification: none; flexible modifications: oxidation on methionine for the primary search, and dioxidation of methionine and glutamine to pyro-glutamine at the refinement stage; protease specificity: tryptic allowing 2 miscleavages for the primary search and semitryptic allowing 5 miscleavages at the refinement stage. Only spectra with loge <-2 were included in the final report. Peptides belonging to target proteins were manually inspected to determine exact processed sequences.

**Limited proteolysis of SARS-CoV-2 nsp7-10 polyprotein by Mpro**

The nsp7-10 cleavage experiment was performed in the assay buffer (20 mM HEPES pH 7.5, 5% glycerol, 1 mM DTT, 100 mM NaCl) at room temperature. 150 μL reaction was set with WT Mpro and nsp7-10 polyprotein in a molar ratio of 1:2. 10 μL of the reaction mix was taken after regular time intervals starting from 0 to 60 min. Using the SDS-PAGE based approach, the stepwise order of the polyprotein proteolysis was analyzed.

**Structural integrative modeling of nsp7-8 using I-TASSER**

For the structural predictions of the nsp7-8 polyprotein, a similar integrative modeling approach to nsp7-11 modeling was employed using the I-TASSER server.

For nsp7-8 modeling, we employed a similar two-run approach using the amino acid sequence and varied the experimental input parameters. Run 1 included using PDB:6YHU as a template, along with distance constraints from crosslinking MS (XL-MS), and secondary structural restraints for nsp7 and nsp8 from solved X-ray crystal structures as guided by HDX-MS to generate Models 1-5 **(Fig. S10B)**.  Run 2 also used PDB:6YHU as a template, distance constraints from crosslinking MS (XL-MS), and secondary structure restraints for nsp8 from the crystal structure as guided by HDX-MS to generate Models 6-10 **(Fig. S10B)**. We employed the secondary structural elements from PDB: 6YHU as the HDX-MS solvent exchange profile of the individual proteins is similar to the nsp7-8 polyprotein **(Fig. 3A).** These data suggest that the individual proteins and polyprotein contain similar secondary structure elements. The 10 output models were assessed against the experimental (i) HDX-MS, (4) XL-MS and the (iii) SAXS data, similar to what we did for nsp7-11 polyprotein.

As the HDX-MS profile of the nsp7-8 polyprotein resembles the profile of nsp7-8 within the nsp7-11 polyprotein **(Fig. 3A)**, we used models of nsp7-8 as a structural template for the structural modeling of nsp7-11 **(Fig. 3B)**. Models 5 and 6 were selected as templates for nsp7-11 modeling based on their agreement with the XL-MS and HDX-MS data. Model 5 is most consistent with the distance restraints obtained by XL-MS, whereas Model 6 is most consistent with the solvent exchange profiles obtained by HDX-MS. As such, we wanted to limit experimental bias and maximize the conformational variety for the integrative modeling of nsp7-11.

The nsp7-8 integrative structures have been deposited in the PDB-Dev databank under accession codes PDBDEV_00000119. They are also provided in the SM as PyMOL sessions.

**Results**

**Structural integrative modeling of SARS-CoV-2 nsp7-8 polyprotein**

Similarly, as with nsp7-11, we first used analytical SEC coupled to a multi-angle light scattering (MALS) and SAXS detection (SEC-MALS-SAXS) to analyze the in-solution assembly state and structural features of the nsp7-8 polyprotein. Two major peaks were detected from the SEC-MALS analysis of the nsp7-8 polyprotein with the main peak corresponding to an equilibrium between monomeric and dimeric states **(Fig. S9A).** SAXS analysis was conducted for both the oligomeric states by using evolving factor analysis (EFA) to separate the scattering of the monomer and dimer components **(Fig. S9B)**. A linear Guinier plot for both the states suggested the presence of stable protein samples with no aggregation **(Fig. S9C)**. A bell-shaped (Gaussian) curve at lower q values was observed in the Kratky plot indicating that the sample contains folded domains with no significant disorder **(Fig. S9D)**. The pair-distance distribution function, P(r), which is related to the shape of the sample, indicated a globular shaped protein for both the monomeric and dimeric forms **(Fig. S9E).** The Rg and the Dmax values calculated from the P(r) are: 33.7 Å and 118 Å for the dimer, and 25.2 Å and 88 Å for the monomeric nsp7-8 **(Table S2).**

The I-TASSER workflow generated an ensemble of ten nsp7-8 models **(Fig. S10B).** All models show the characteristic nsp7 helical bundle with a more variable conformation of nsp8. Unlike nsp7-11, the nsp7-8 models don't adopt the “golf-club” conformation in the nsp8 domain.  Analysis of the nsp7-8 polyprotein models suggest that the nsp8 domain is much more globular in the context of this polyprotein intermediate. Visual inspection of models in nsp7-11 Group A suggest that this extended helical conformation is stabilized by nsp7 on one side and by nsp9 and nsp10 from the other. Thus, in the case of nsp7-8, the lack of scaffolding by the absent nsp9-11 domains may prevent the extended helical conformation. This absence of extended nsp8 helix in the context of nsp7-8 might enhance nsp7-8 polyprotein stability, limiting its ability to sample dynamic conformations and ultimately making it less amenable for cleavage by Mpro.

All the models satisfied most of the crosslinks with distances equal to or less than 30 Å, the theoretical upper limit distance for the DSSO crosslinker **(Fig. S10C)**, with Model 7 able to satisfy all of the crosslinks. Moreover, all crosslinks are satisfied across all the models.

Next, we mapped the percent deuterium uptake at 10 s incubation in deuterated buffer on the models to assess the agreement of the modeled secondary structure elements to the in-solution dynamics of the polyprotein **(Fig. S10D)**. Using HDXer software we calculated deuterated fractions for peptide segments corresponding to the experimental data as a function of the experimental deuterium exposure times. We then plotted the computationally derived 10 s percent deuterium value for each model versus the experimentally determined 10 s percent deuterium value and calculated the RMSE. We observed similar agreement of all models with the experimental HDX-MS data, with only ~0.1 RMSE difference between Model 7, which shows highest RMSE (least agreement), and Model 1, which shows the lowest RMSE (best agreement). Interestingly, the average RMSE for the individual domains was greater for nsp7 compared to nsp8, suggesting that the models do a better job at modeling nsp8. Model 6 was the only model where nsp7 had better agreement with HDX-MS experimental data compared to nsp8. For this reason, Model 6 was chosen as one of the template models for the nsp7-11 modeling.

Next, we reconstructed a three-dimensional shape (bead model) for the monomeric nsp7-8. Based on evaluation of the bead models **(Table S2)**, the reconstruction seemed somewhat unstable, which may be caused by conformational flexibility in solution. Nonetheless, all the models fitted well in the envelope **(Fig. S8E)**. The theoretical scattering profile of each monomer model was calculated using the *CRYSOL* software package and fitted against the experimental scattering profile **(Fig. S8F)**. The calculated scattering profiles fit well to the experimental scattering profile, as seen with the low χ^2^ values. The calculated radius of gyration (R_g_) values for the models were also comparable with the experimental R_g_ value of the monomeric nsp7-8 polyprotein (experimental R_g_ is 25.2 Å) calculated from the P(r) function, suggesting that the overall size of the model matches with the actual size of the polyprotein in solution.

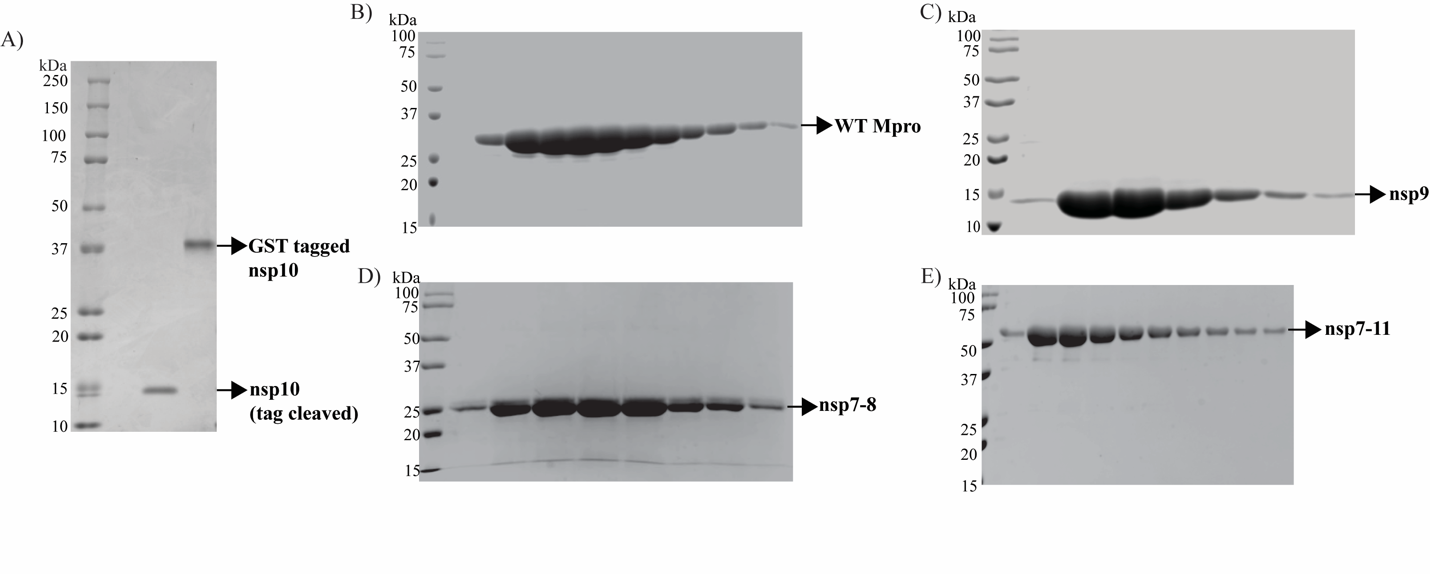

**Fig. S1. Confirmation of protein expression and purification.** SDS-PAGE of purified (A) nsp10, (B) WT Mpro, (C) nsp9, (D) nsp7-8, and (E) nsp7-11.

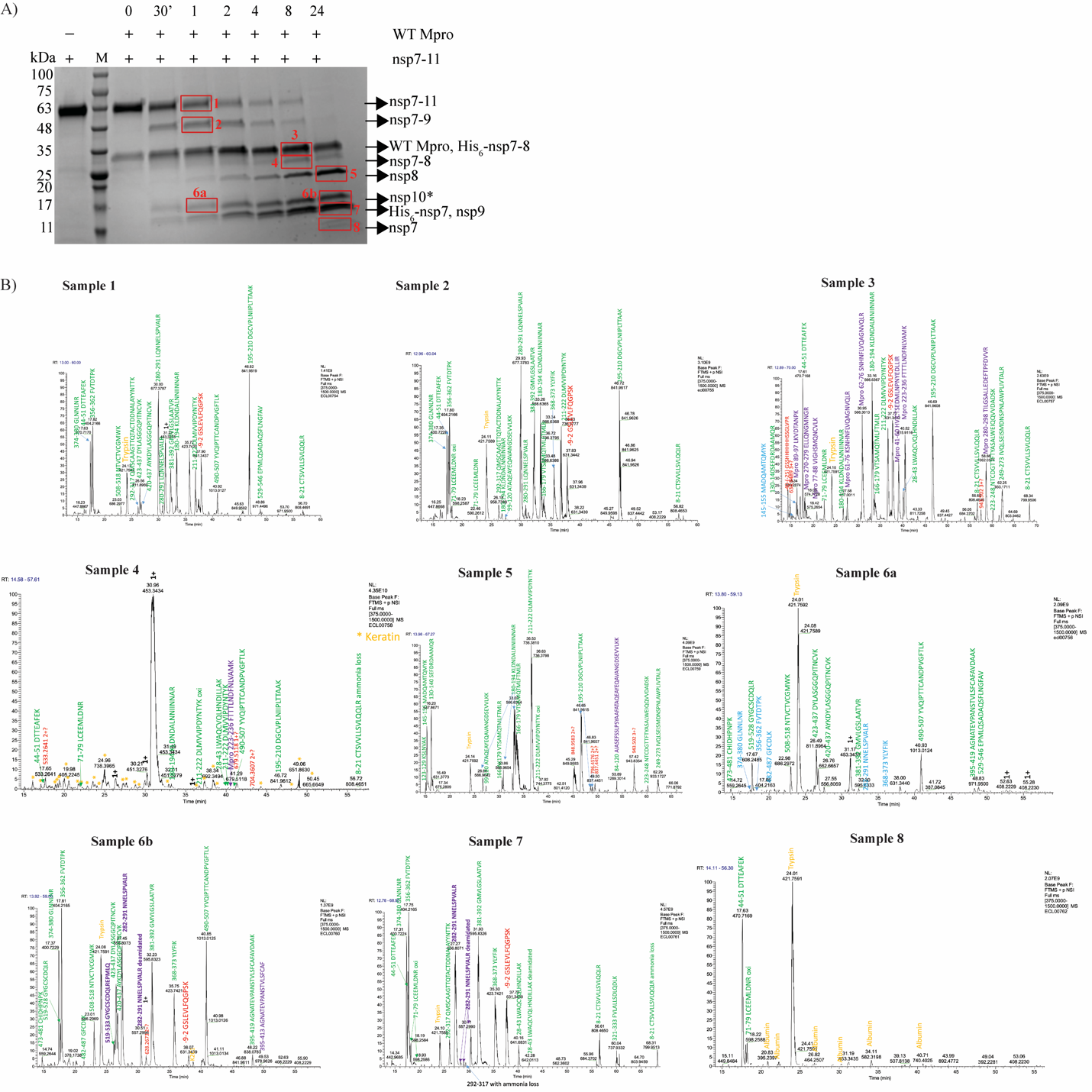

**Fig. S2. Validation of nsp7-11 *in vitro* processing by WT Mpro.** (A) SDS-PAGE of nsp7-11 processed by WT Mpro with representative protein bands confirmed by LC-MS/MS marked in red. *After 30 mins of incubation, the gel band corresponds to nsp10-11 polyprotein intermediate and after 24 h of incubation, the band corresponds to nsp10. The molecular mass of nsp11 is only 1.3 kDa so it couldn’t be resolved on the SDS-PAGE gel. (B) Sample traces of peptide IDs in LC-MS/MS of the marked protein bands.

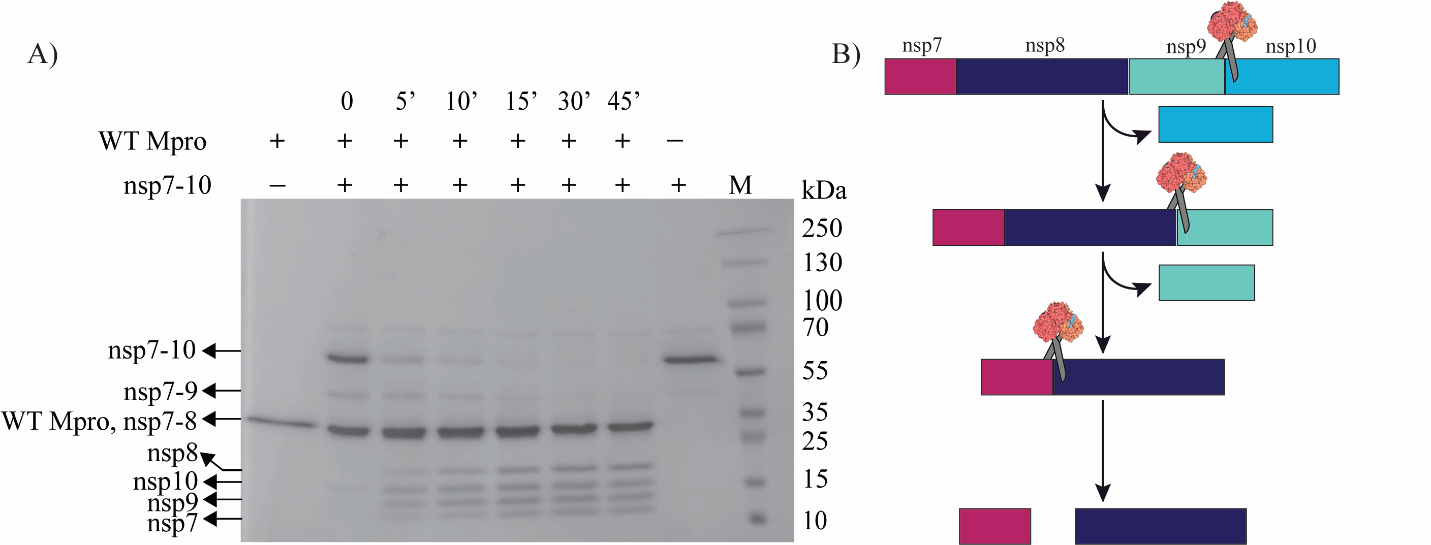

**Fig. S3. *In vitro* analysis of SARS-CoV-2 nsp7-10 polyprotein processing by WT Mpro.** (A) SDS-PAGE of nsp7-10 processed by WT Mpro. (B) Schematic representation of the cleavage order of the nsp7-10 polyprotein by WT Mpro.

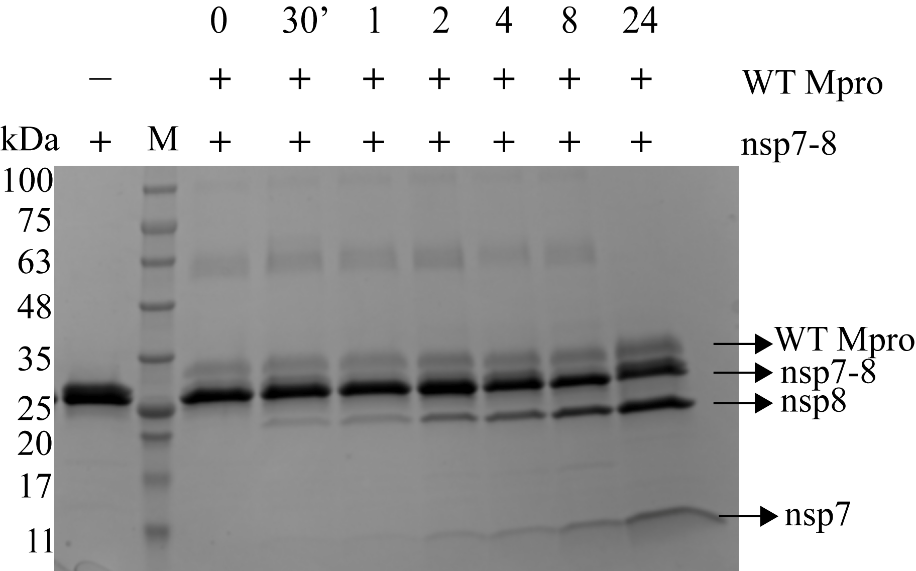

**Fig. S4.** ***In vitro* analysis of nsp7-8 polyprotein processing by WT Mpro.** SDS-PAGE of nsp7-8 processed by WT Mpro.

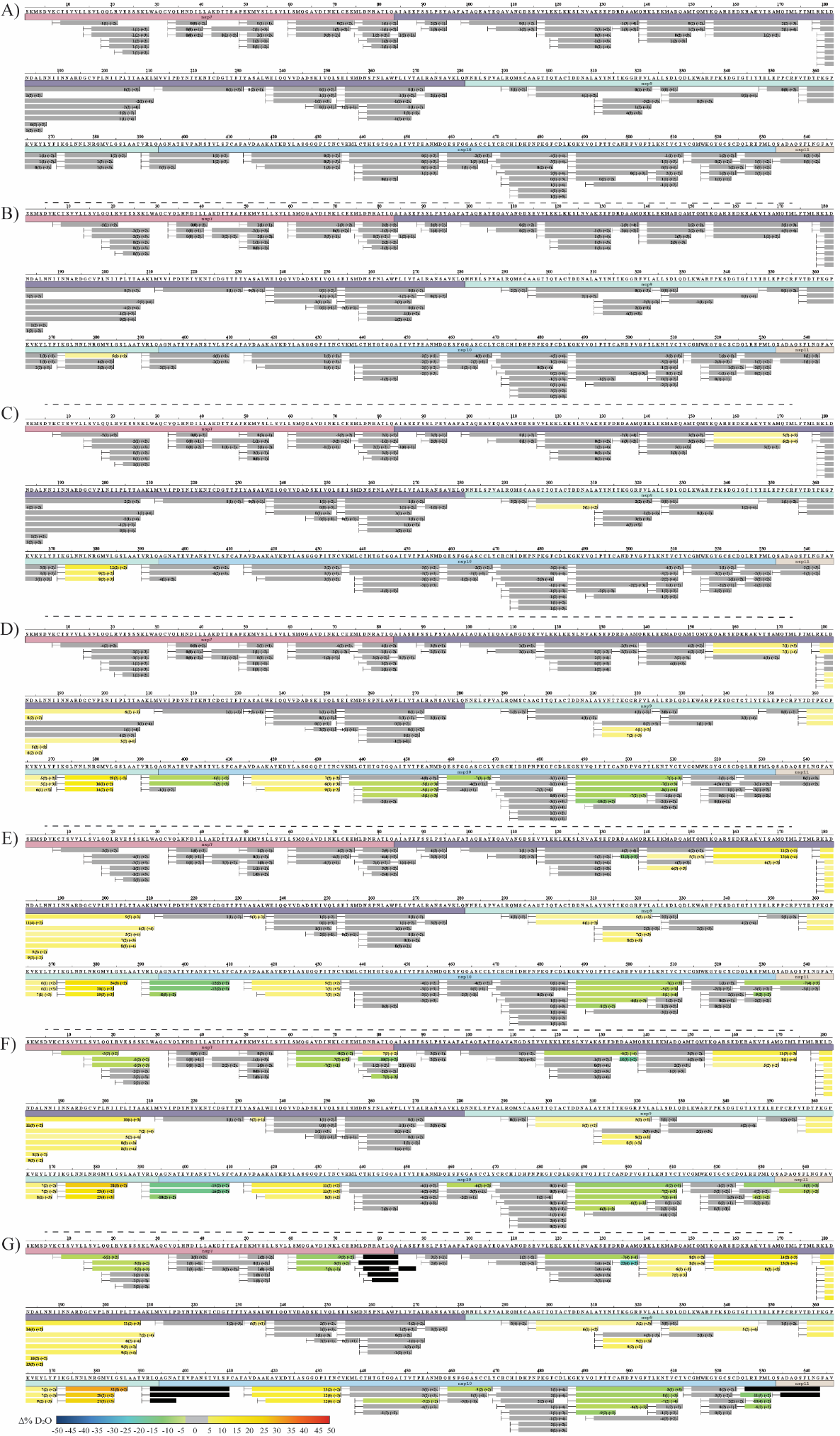

**Fig. S5.** **Pulsed HDX-MS of nsp7-11 with WT Mpro.** Sequence coverage of differential pulsed HDX-MS analysis of nsp7-11 versus nsp7-11 with WT Mpro after (A) 600 s, (B) 1800 s, (C) 3600 s, (D) 7200 s, (E) 14400 s, (F) 28800 s, and (G) 86400 s of cleavage reaction time. Color scale represents changes in deuterium uptake (30 s incubation in deuterated buffer) over the course of the cleavage reaction, with gray representing no significant change in deuterium uptake and black representing peptides no longer identifiable in the mass spectrometer.

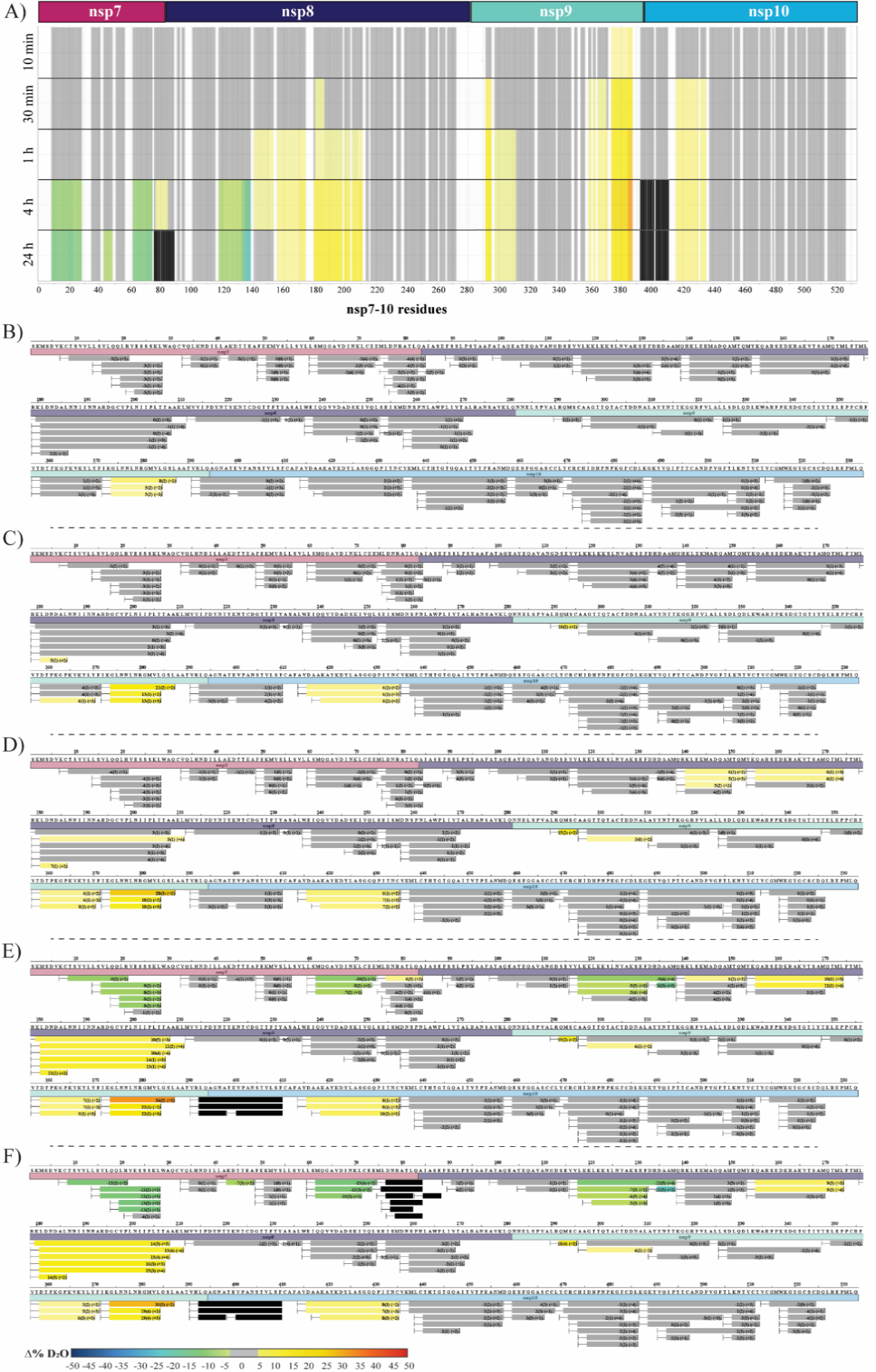

**Fig S6.** **Pulsed HDX-MS of nsp7-10 with WT Mpro.** (A) Pulsed HDX-MS analysis of nsp7-10 with WT Mpro. Sequence coverage of differential pulsed HDX-MS analysis of nsp7-11 versus nsp7-11 with WT Mpro after (B) 600 s, (C) 1800 s, (D) 3600 s, (E) 14400 s, and (F) 86400 s of cleavage reaction time. Color scale represents changes in deuterium uptake (30 s incubation in deuterated buffer) over the course of the cleavage reaction, with gray representing no significant change in deuterium uptake, white denoting no sequence coverage, and black representing peptides no longer identifiable in the mass spectrometer.

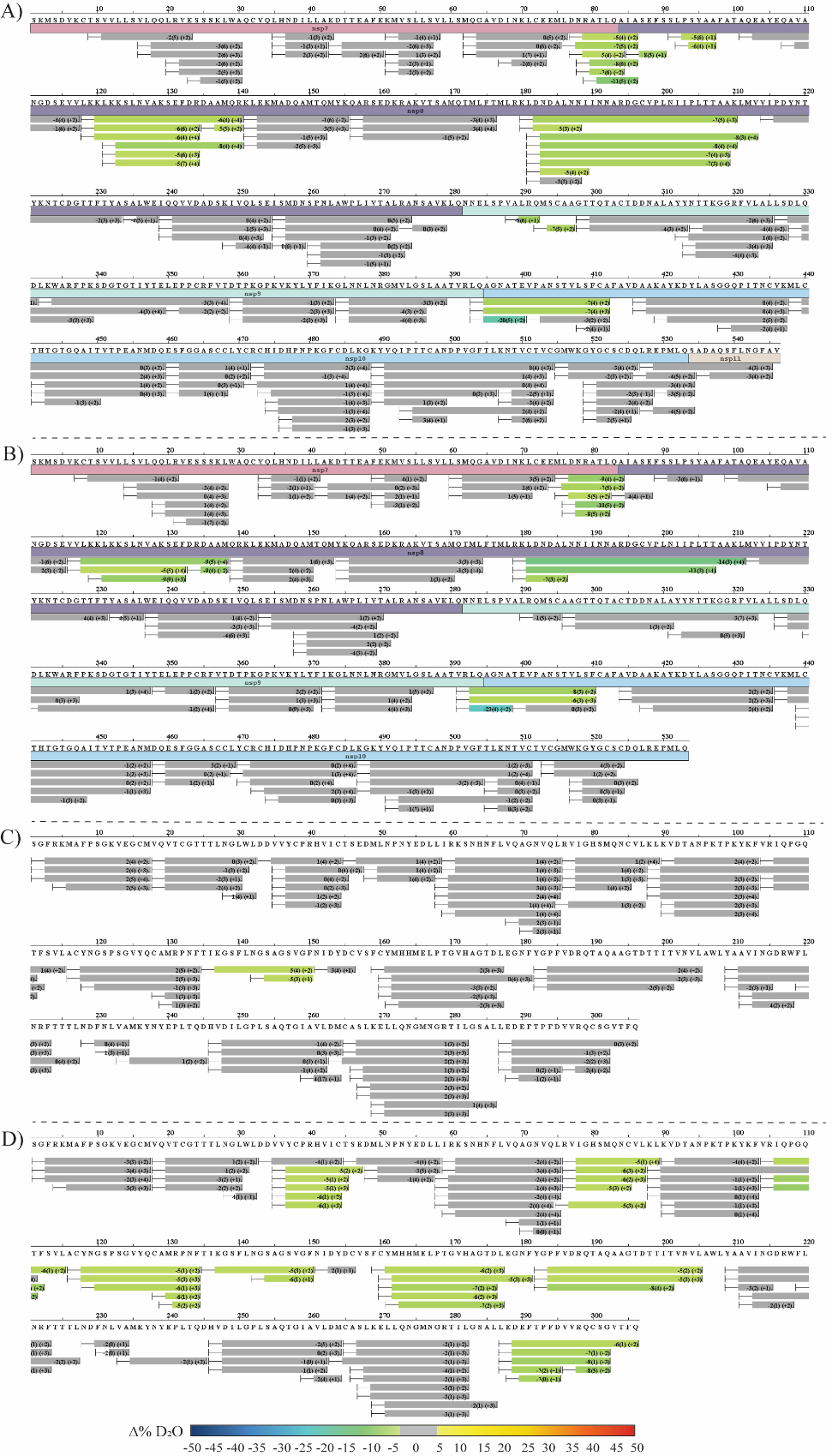

**Fig. S7. HDX-MS of C145A Mpro nsp7-10 complex.** Sequence coverage of differential HDX-MS analysis of (A) nsp7-11 versus nsp7-11 with C145A Mpro, (B) nsp7-10 versus nsp7-10 with C145A Mpro, (C) C145A Mpro versus C145A Mpro with nsp7-11 at 10 s, 60 s, 5 min, 30 min, 1 h, and 12 h incubation in deuterated buffer, and (D) C145A Mpro versus C145A Mpro with nsp7-11 at 12 h incubation in deuterated buffer only. Color scale represents changes in deuterium uptake, with gray representing no significant change in deuterium uptake.

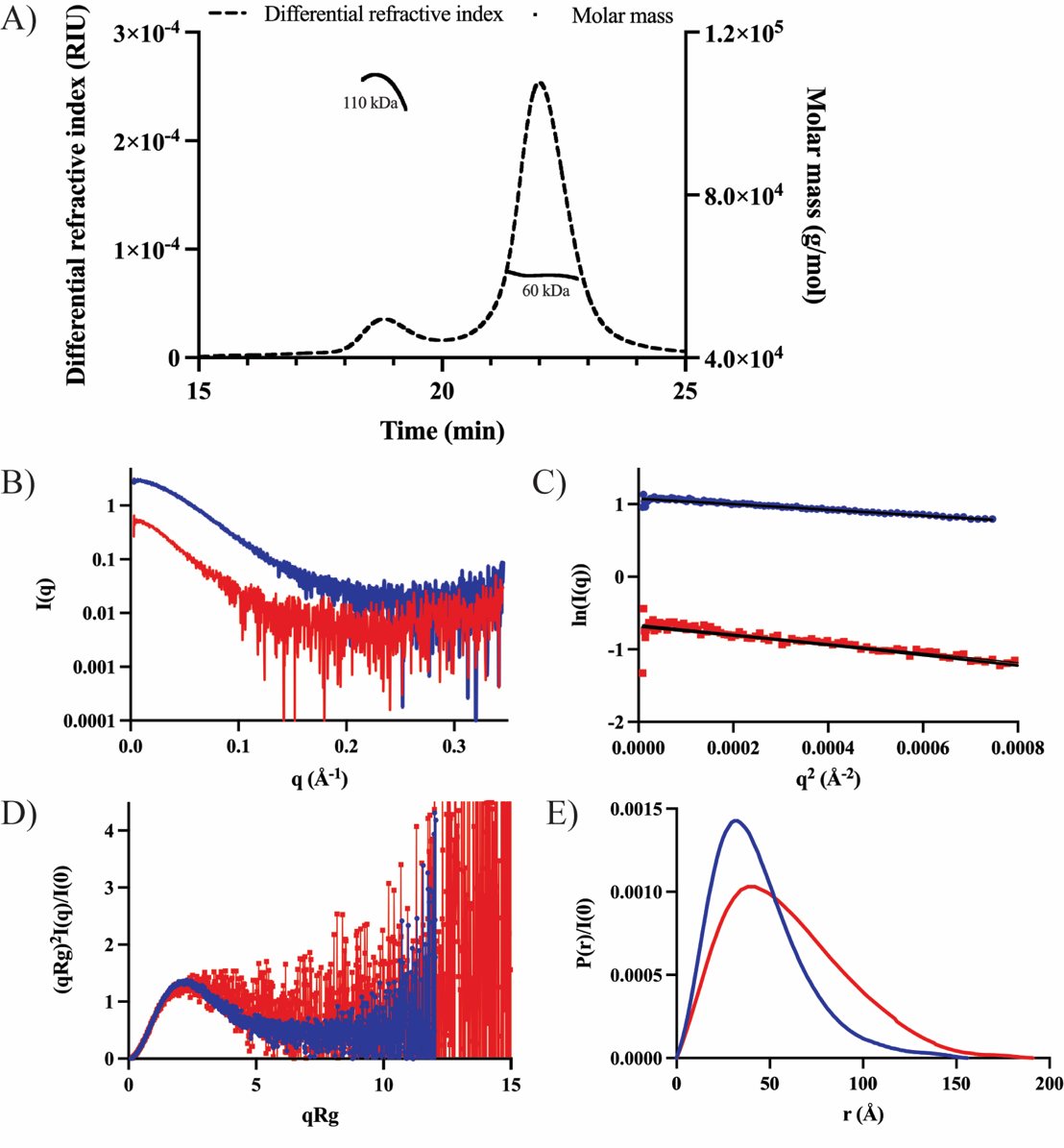
**Fig. S8. SEC-MALS-SAXS analysis of nsp7-11 polyprotein.** A) SEC-MALS chromatogram of nsp7-11 polyprotein. Dashed line represents the DRI (differential refractive index) signal, and the plain line represents the molecular weight of the corresponding peaks. B) SAXS scattering profiles with I(q) versus q as log-linear plots, C) Guinier fit, D) Dimensionless Kratky plots and E) Pair-wise distribution curve of monomeric (blue) and dimeric (red) nsp7-11 polyprotein.

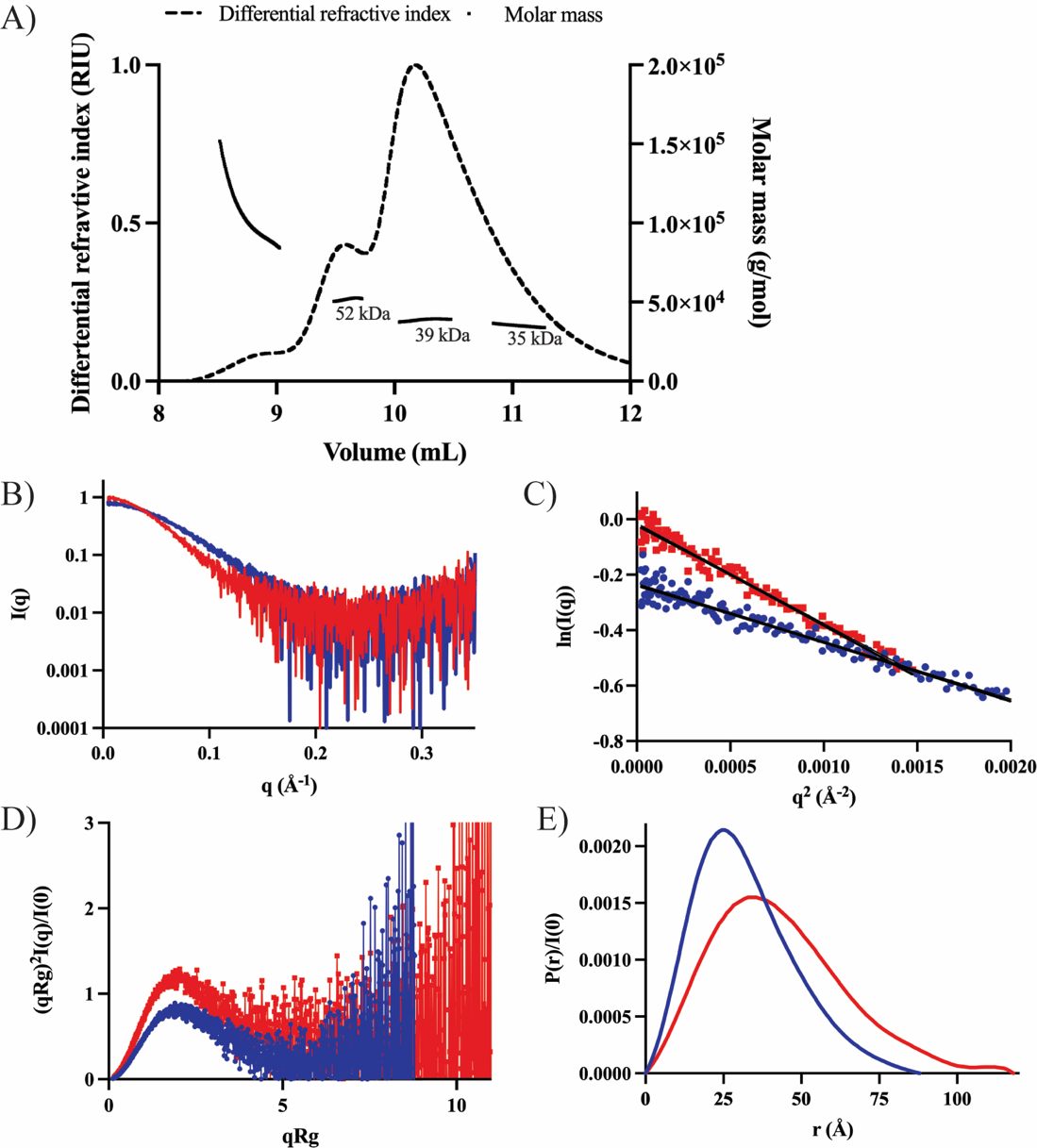

**Fig. S9. SEC-MALS-SAXS analysis of nsp7-8 polyprotein.** A) SEC-MALS chromatogram of nsp7-8 polyprotein. Dashed line represents the DRI (differential refractive index) signal, and the plain line represents the molecular weight of the corresponding peaks. B) SAXS scattering profiles with I(q) versus q as log-linear plots, C) Guinier fit, D) Dimensionless Kratky plots and E) Pair-wise distribution curve of monomeric (blue) and dimeric (red) nsp7-8 polyprotein.

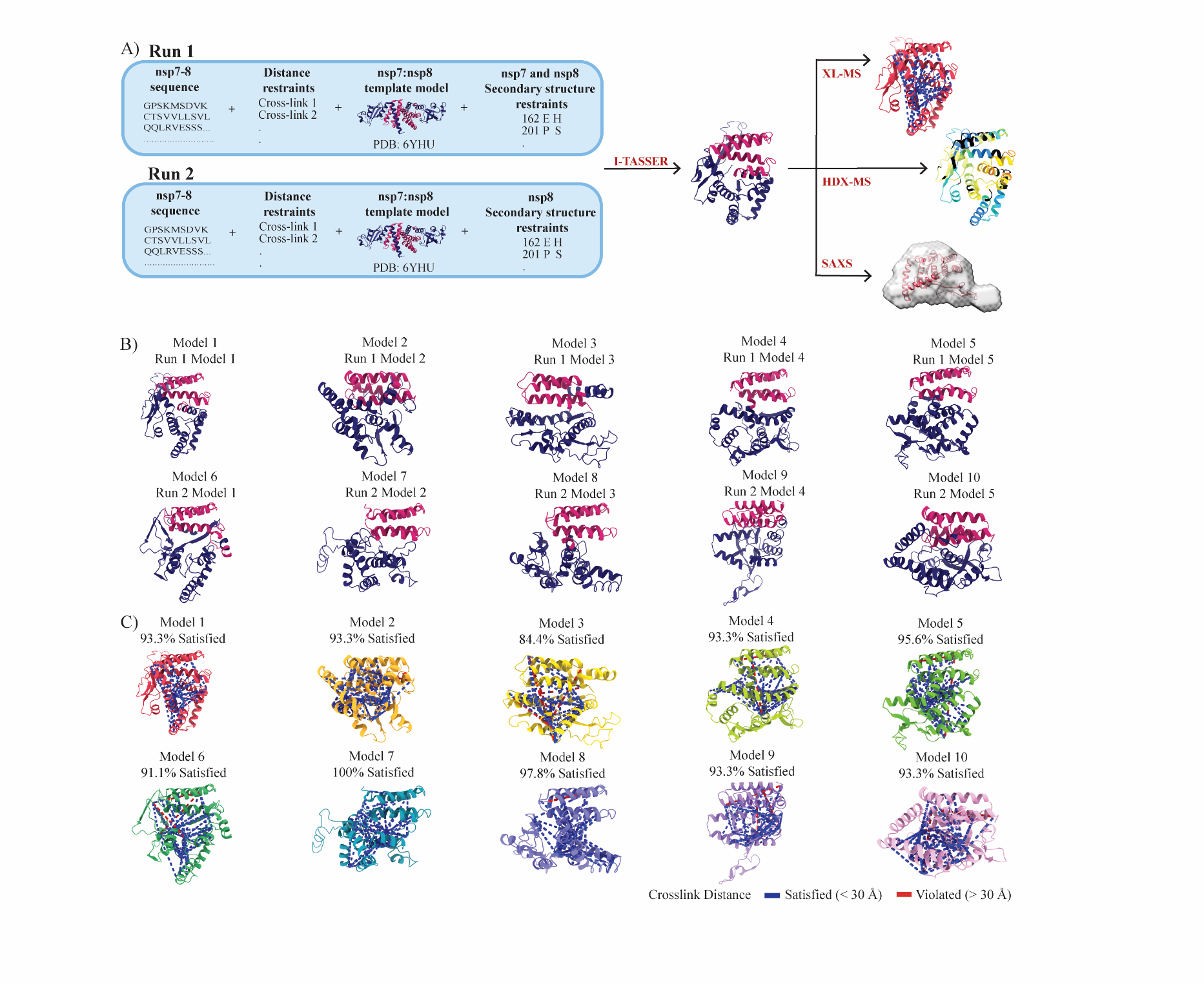

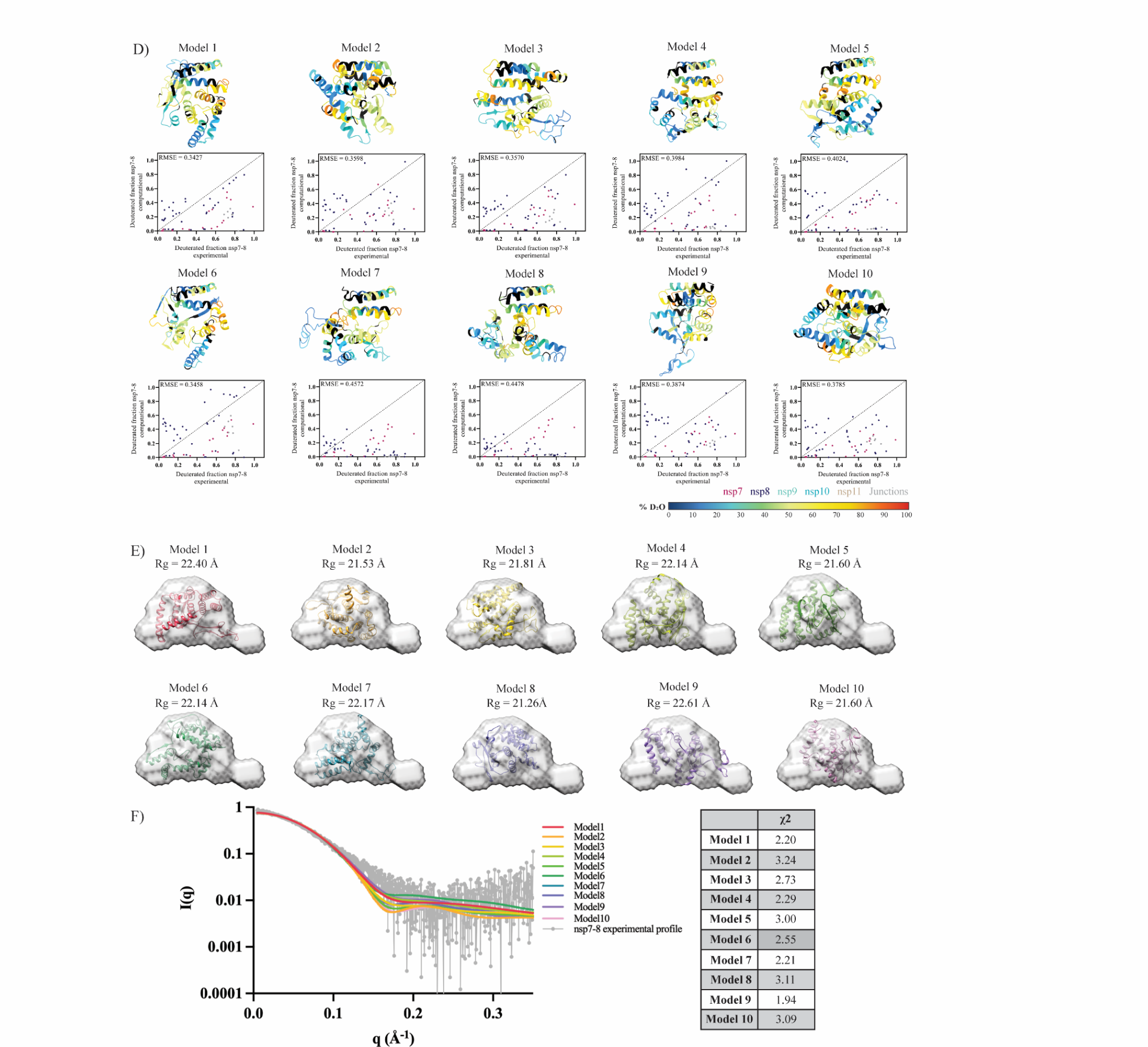

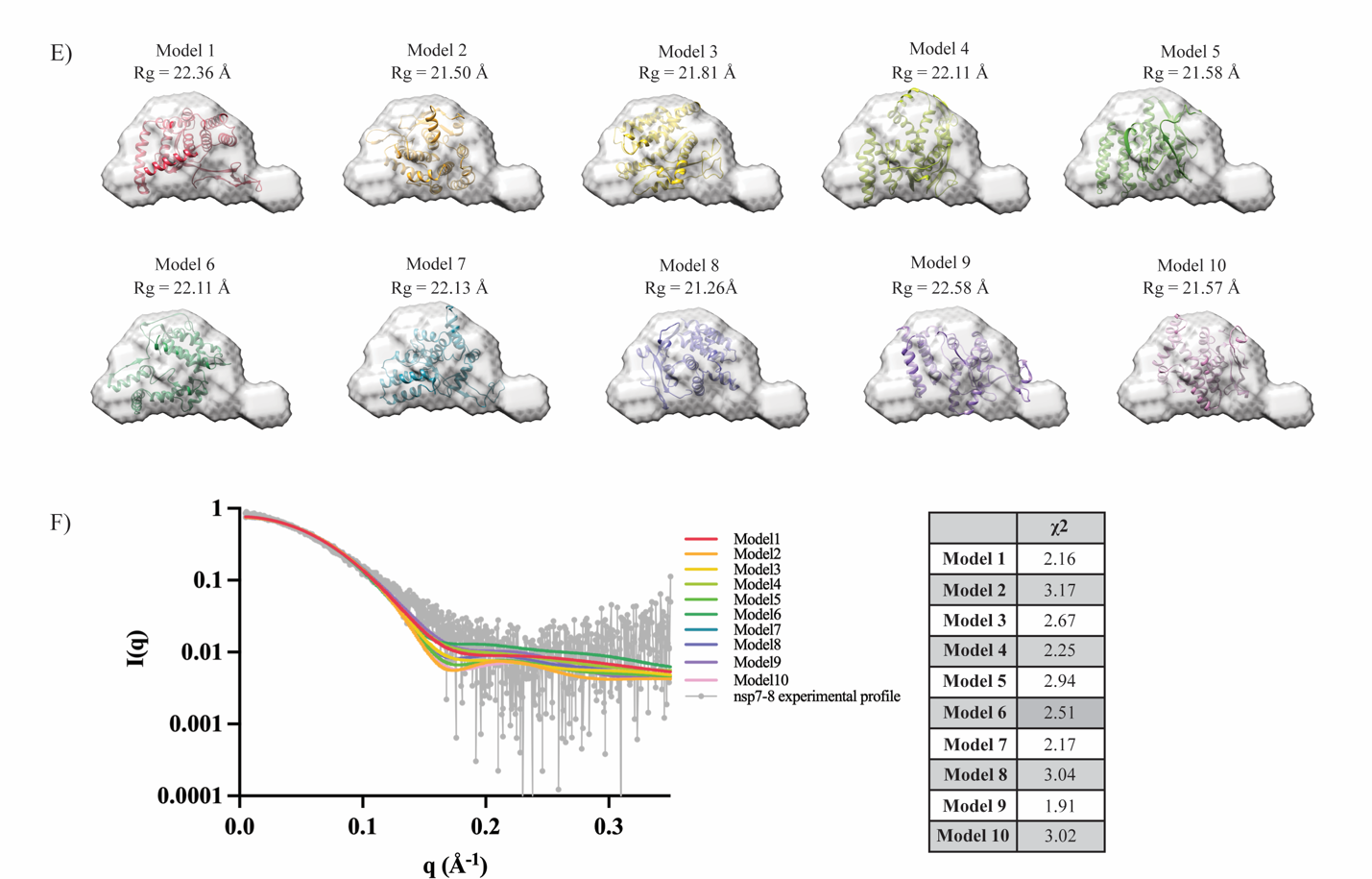

**Fig. S10. Integrative structural modeling approach and assessment of nsp7-8 top ten models.** (A) Scheme of integrative structural modeling workflow for nsp7-8 polyprotein. One model is shown to represent all ten generated. (B) Top ten nsp7-8 models with nsp7 colored in magenta and nsp8 in dark purple. (C) Mapping nsp7-8 intra-protein crosslinks onto all nsp7-8 models. Satisfied crosslinks equal to or less than 30 Å are shown in blue and violated crosslinks greater than 30 Å are shown in red. Percent of crosslinks satisfied is reported. (D) Top ten nsp7-8 models are colored based on percent deuterium value at 10 s incubation in deuterated buffer. Black indicates no sequence coverage in the HDX-MS profiling experiment. Plots of calculated 10 s deuterated fraction versus experimental deuterated fraction shown under each corresponding model with agreement score reported as the RMSE value. (E) Fitting of all nsp7-8 models into the reconstructed SAXS envelope and Rg values are reported under the model. (F) Theoretical scattering profile of all nsp7-8 models fitted against the experimental profile with **χ**2 values of the fit reported.

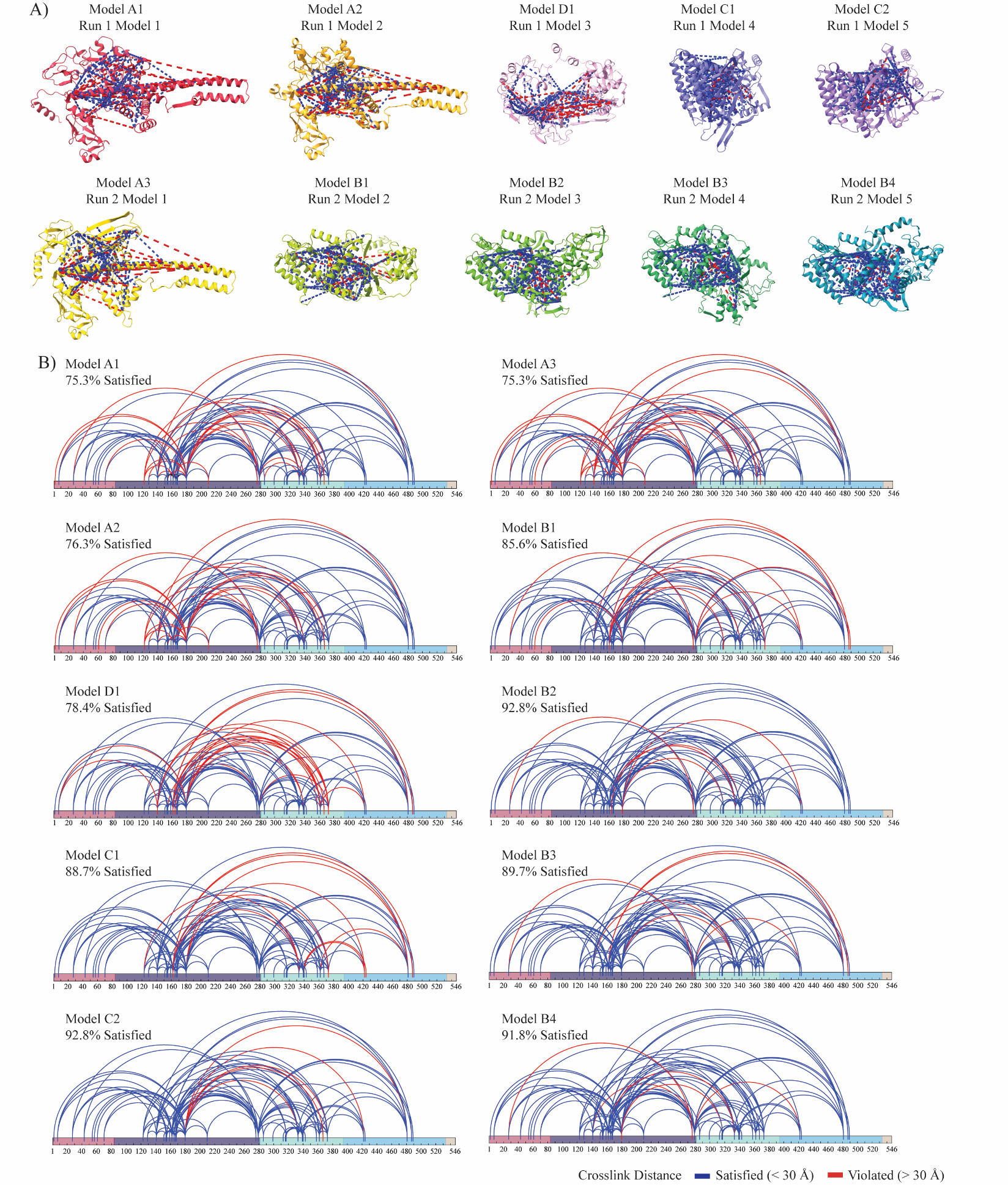

**Fig. S11. Assessment of nsp7-11 top ten models based on XL-MS data.** (A) Top ten nsp7-11 models with all nsp7-11 intra-protein crosslinks mapped. Satisfied crosslinks equal to or less than 30 Å are shown in blue and violated crosslinks greater than 30 Å are shown in red. (B) Alternative view of all nsp7-11 intra-protein crosslinks mapped onto nsp7-11 sequence with nsp7 in magenta, nsp8 in purple, nsp9 in teal, nsp10 in cyan and nsp11 in tan. Percent of crosslinks satisfied is reported for each model.

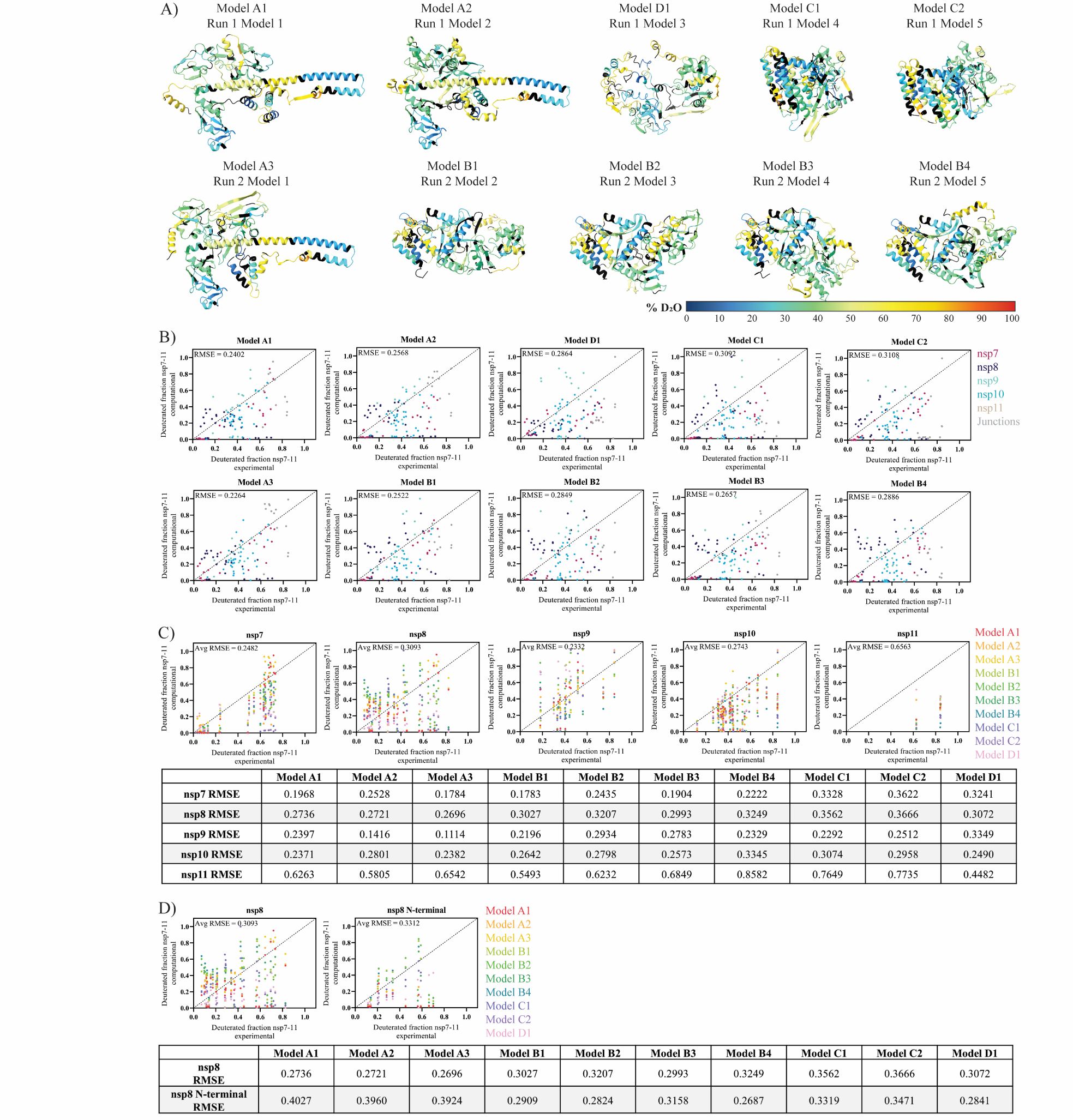

**Fig. S12. Assessment of nsp7-11 top ten models based on HDX-MS data.**  (A) Top ten nsp7-11 models colored based on percent deuterium value after 10 s incubation in deuterated buffer. Black indicates no sequence coverage in the HDX-MS experiment. (B) Plots of calculated 10 s deuterated fraction for each model versus experimental deuterated fraction with agreement score reported as the RMSE value. Data points are colored based on peptide sequence with nsp7 in magenta, nsp8 in purple, nsp9 in teal, nsp10 in cyan, nsp11 in tan, and junctions in gray. (C) Plots of calculated 10 s deuterated fraction for each nsp versus experimental deuterated fraction. Data points are colored based on models. The RMSE value per each individual nsp for each model is reported in the table below. The average agreement scores of given individual nsp for all models is reported as the average RMSE value on plot. (D) Comparison of agreement of nsp8 and nsp8 N-terminal (residues 92-213) only to experimental HDX-MS data. Plots showing calculated 10 s deuterated fraction versus experimental deuterated fraction with data points are colored based on models. The RMSE value for each model is reported in the table below and the average agreement score for all models is reported as the average RMSE value on plot.

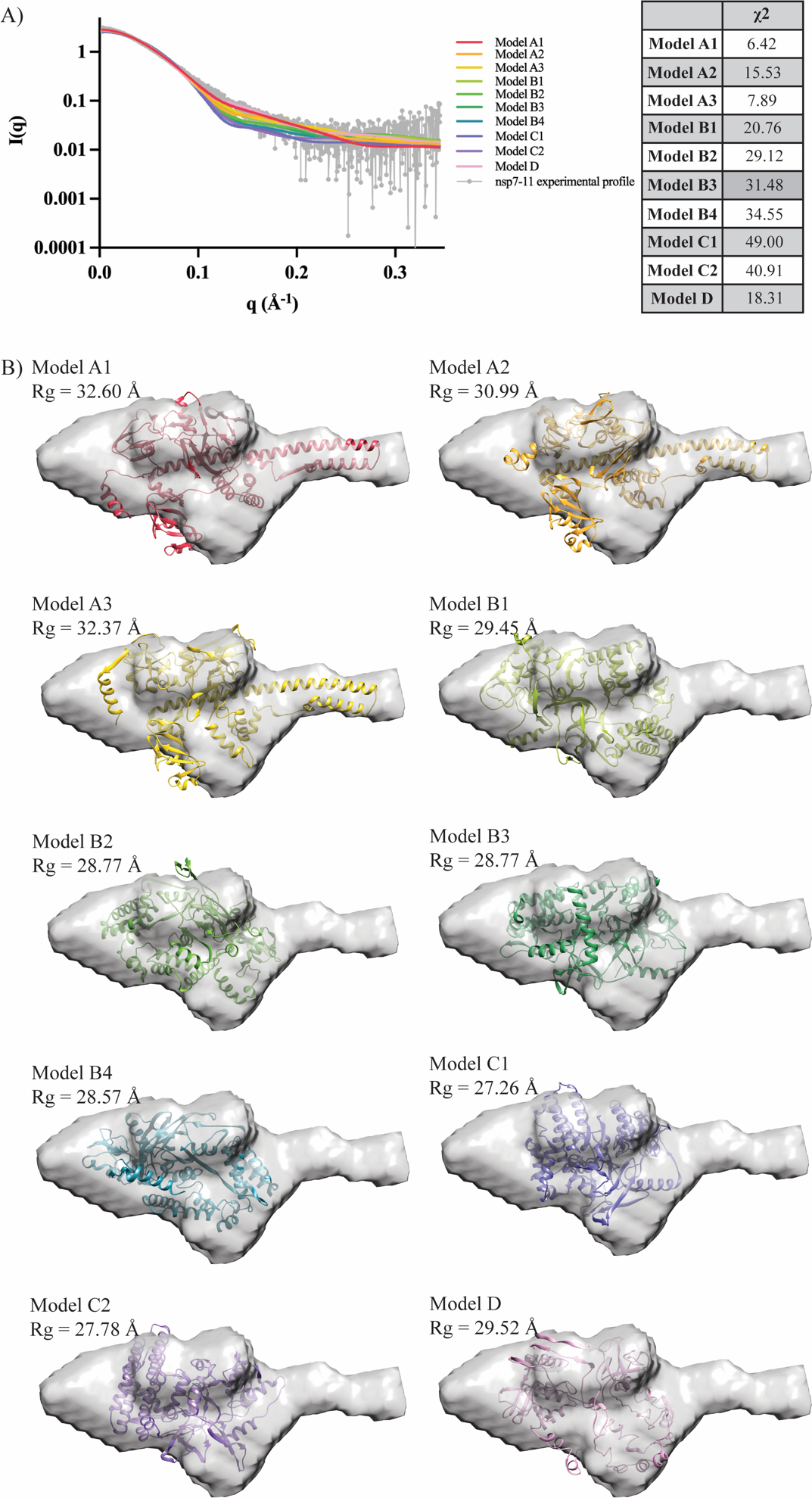

**Fig. S13. Assessment of nsp7-11 top ten models based on SAXS.** (A) Theoretical scattering profile of top 10 models fitted against the experimental profile with χ2 values. (B) All models fitted in the reconstructed SAXS envelopes with Rg value reported for each model. The theoretical Rg value for the models and the χ2 of the fit are calculated using *CRYSOL*.

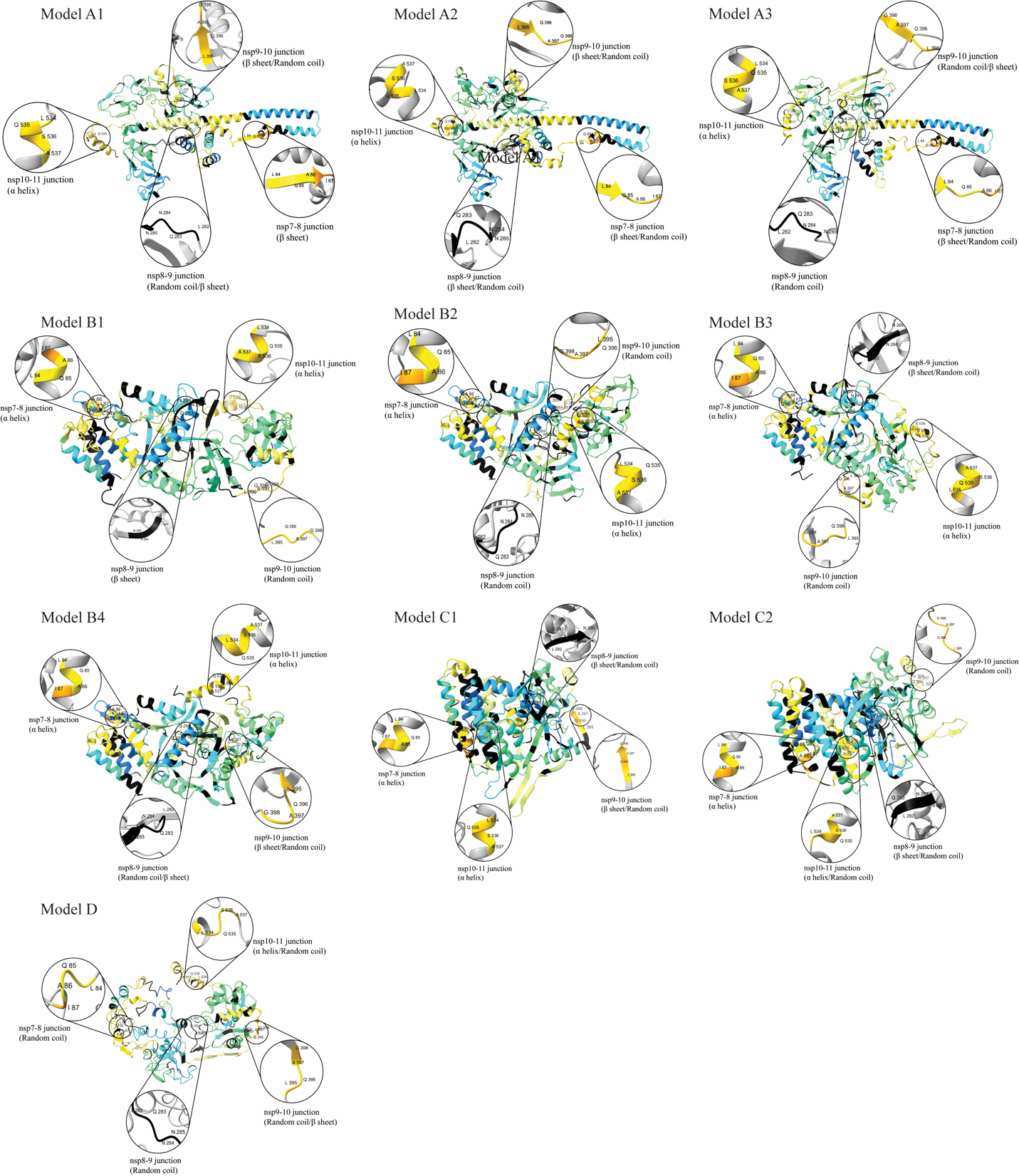
**Fig. S14. Assessment of the junction sites of top ten models of nsp7-11.** Analysis of the secondary structure elements of the junction sites in all nsp7-11 models. Models are overlaid with percent deuterium uptake values from **Fig. S12A**.

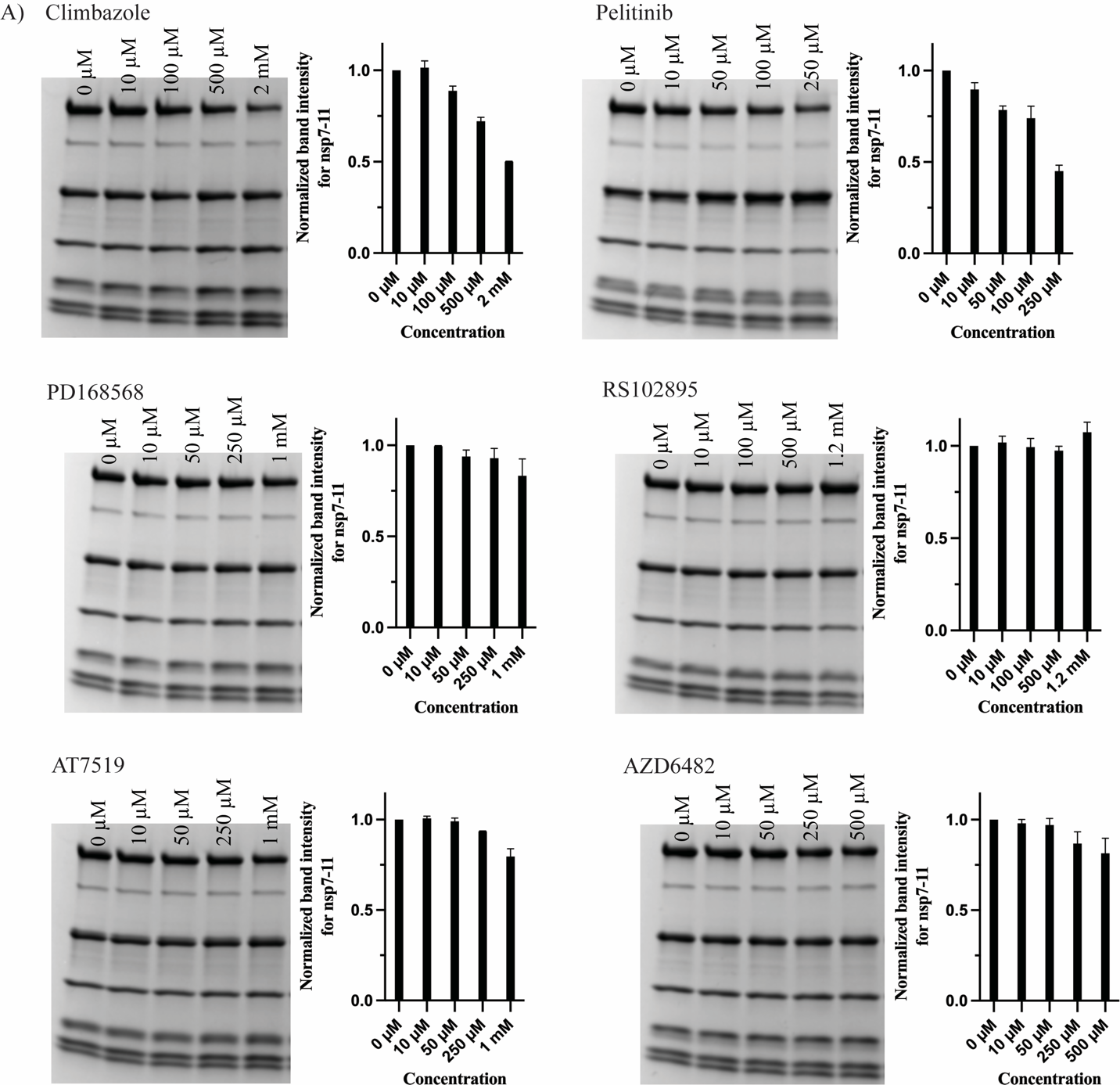

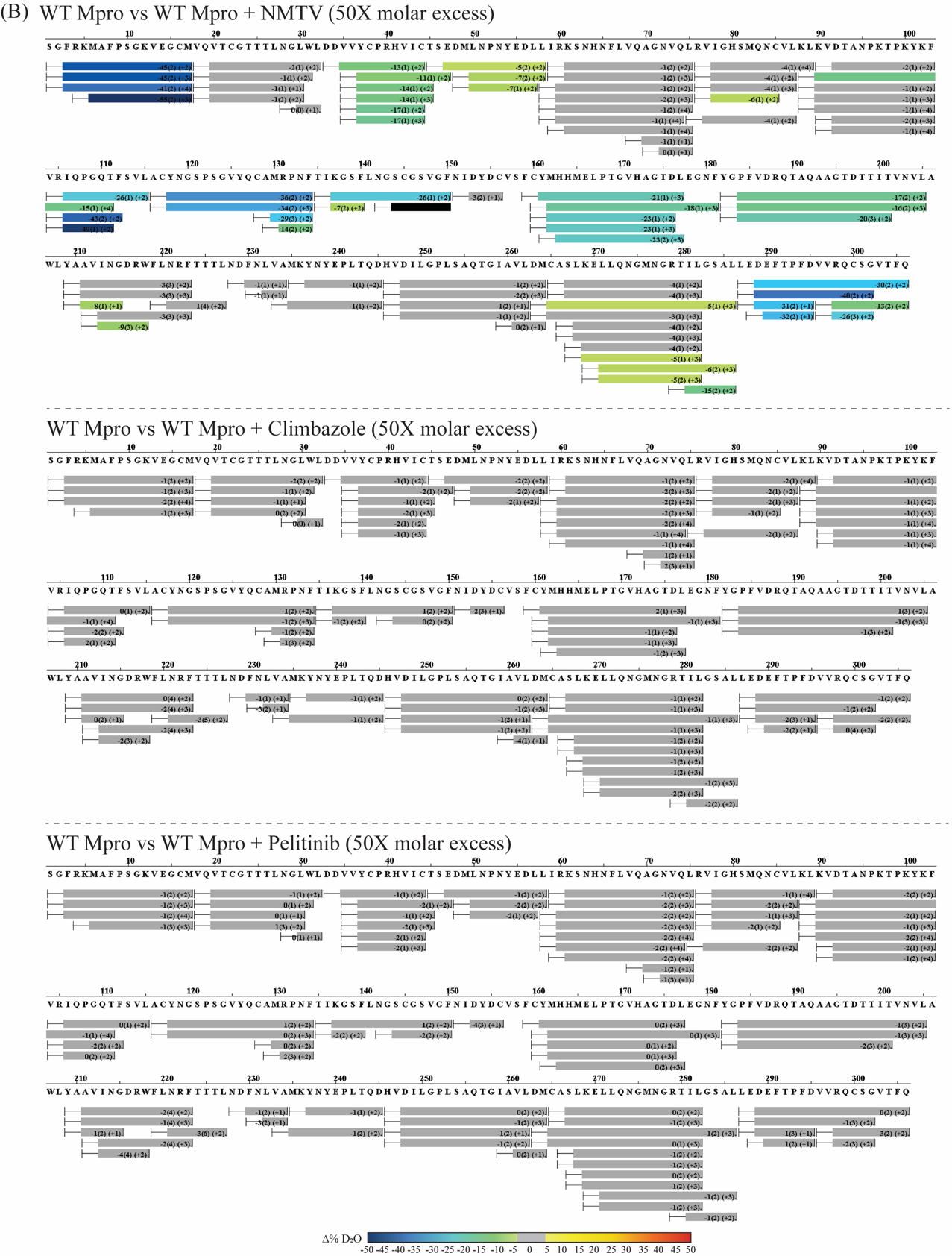

**Fig. S15. Analysis of Mpro crystallographic binders and NMTV on nsp7-11 polyprotein processing by WT Mpro.** (A) SDS-PAGE gels of nsp7-11 polyprotein processing by WT Mpro in the presence of small molecule binders after 24 h. The band intensities were calculated for the nsp7-11 gel band using ImageJ and plotted against the concentration of the binders. The identity of the substrate, intermediate, and final cleavage product gel bands can be referenced to **Fig. S2**. (B) Sequence coverage of differential HDX-MS analysis of WT Mpro versus WT Mpro with binders at 12 h incubation in deuterated buffer only. Color scale represents changes in deuterium uptake with gray representing no significant change in deuterium uptake.

**Table S1.** HDX-MS experimental conditions and data analysis parameters from the guidelines of the IC-HDX-MS community.

| **Data Set (Figure 1C, S5)** | **nsp7-11** | | | | | | | | **nsp7-11 : WT Mpro** | | | | | |
| --- | --- | --- | --- | --- | --- | --- | --- | --- | --- | --- | --- | --- | --- | --- |
| Reaction details | 50 mM HEPES, 500 mM NaCl, 1 mM TCEP, pH 8.0, 4 °C | | | | | | | | | | | | | |
|  | 600 s | 1800 s | | | 3600 s | | 7200 s | | | 14400 s | | 28800 s | | 86400 s |
| Deuterium labeling time (sec) | 30 | | | | | | | | | | | | | |
| HDX quench reaction details | 5 M urea, 1% TFA, pH = 2.0, 4 °C | | | | | | | | | | | | | |
| Back-exchange | estimated from input recovery estimate of 0.7 and deuterated buffer concentration of 0.8 | | | | | | | | | | | | | |
| # of peptides | 119 | | | | | | | | | | | | | |
| Sequence coverage | 99% | | | | | | | | | | | | | |
| Average peptide length / Redundancy | 13 / 3.314 | | | | | | | | | | | | | |
| Replicates (biological or technical) | 3 (technical) | | | | | | | | | | | | | |
| Repeatability (average STD) | 1.478 | | 1.811 | | 1.683 | | 1.460 | | | 1.984 | | 2.038 | | 2.163 |
| Significant differences in HDX | > 5% D (unpaired t-tests at each time point, p-value < 0.01) | | | | | | | | | | | | | |
| **Data Set (Figure S6)** | **nsp7-10** | | | | | | | | **nsp7-10 : WT Mpro** | | | | | |
| Reaction details | 50 mM HEPES, 500 mM NaCl, 1 mM TCEP, pH 8.0, 4 °C | | | | | | | | | | | | | |
|  | 600 s | | | 1800 s | | 3600 s | | | | | 14400 s | | 86400 s | |
| Deuterium labeling time (sec) | 30 | | | | | | | | | | | | | |
| HDX quench reaction details | 5 M urea, 1% TFA, pH = 2.0, 4 °C | | | | | | | | | | | | | |
| Back-exchange | estimated from input recovery estimate of 0.7 and deuterated buffer concentration of 0.8 | | | | | | | | | | | | | |
| # of peptides | 107 | | | | | | | | | | | | | |
| Sequence coverage | 99% | | | | | | | | | | | | | |
| Average peptide length / Redundancy | 10 / 3.110 | | | | | | | | | | | | | |
| Replicates (biological or technical) | 3 (technical) | | | | | | | | | | | | | |
| Repeatability (average STD) | 1.723 | | | 1.563 | | 1.743 | | | | | 1.722 | | 2.511 | |
| Significant differences in HDX | > 5% D (unpaired t-tests at each time point, p-value < 0.01) | | | | | | | | | | | | | |
| **Data Set (Figure 2A, S7A)** | **nsp7-11** | | | | | | | | **nsp7-11 : C145A Mpro** | | | | | |
| Reaction details | 50 mM HEPES, 500 mM NaCl, 1 mM TCEP, pH 8.0, 4 °C | | | | | | | | | | | | | |
|  | 30 min | | | | | | | | | | | | | |
| Deuterium labeling time (sec) | 10, 30, 60, 900, 3600 | | | | | | | | | | | | | |
| HDX quench reaction details | 5 M urea, 1% TFA, pH = 2.0, 4 °C | | | | | | | | | | | | | |
| Back-exchange | estimated from input recovery estimate of 0.7 and deuterated buffer concentration of 0.8 | | | | | | | | | | | | | |
| # of peptides | 133 | | | | | | | | | | | | | |
| Sequence coverage | 93% | | | | | | | | | | | | | |
| Average peptide length / Redundancy | 4 / 3.604 | | | | | | | | | | | | | |
| Replicates (biological or technical) | 3 (technical) | | | | | | | | | | | | | |
| Repeatability | 4.178 (average STD) | | | | | | | | | | | | | |
| Significant differences in HDX | > 5% D (unpaired t-tests at each time point, p-value < 0.01) | | | | | | | | | | | | | |
| **Data Set (Figure 2B, 2F, S7B)** | **nsp7-10** | | | | | | | | **nsp7-10 : C145A Mpro** | | | | | |
| Reaction details | 50 mM HEPES, 500 mM NaCl, 1 mM TCEP, pH 8.0, 4 °C | | | | | | | | | | | | | |
|  | 30 min | | | | | | | | | | | | | |
| Deuterium labeling time (sec) | 10, 30, 60, 900, 3600 | | | | | | | | | | | | | |
| HDX quench reaction details | 5 M urea, 1% TFA, pH = 2.0, 4 °C | | | | | | | | | | | | | |
| Back-exchange | estimated from input recovery estimate of 0.7 and deuterated buffer concentration of 0.8 | | | | | | | | | | | | | |
| # of peptides | 97 | | | | | | | | | | | | | |
| Sequence coverage | 90% | | | | | | | | | | | | | |
| Average peptide length / Redundancy | 9 / 2.842 | | | | | | | | | | | | | |
| Replicates (biological or technical) | 3 (technical) | | | | | | | | | | | | | |
| Repeatability (average STD) | 3.454 | | | | | | | | | | | | | |
| Significant differences in HDX | > 5% D (unpaired t-tests at each time point, p-value < 0.01) | | | | | | | | | | | | | |
| **Data Set (Figure 2C-D, 2E-F, 6A, S7C-D)** | **C145A Mpro** | | | | | | | **C145A Mpro : nsp7-11** | | | | | | |
| Reaction details | 50 mM HEPES, 500 mM NaCl, 1 mM TCEP, pH 8.0, 4 °C | | | | | | | | | | | | | |
|  | 30 min | | | | | | | | | | | | | |
| Deuterium labeling time (sec) | 10, 30, 60, 900, 3600, 43200 | | | | | | | | | | | | | |
| HDX quench reaction details | 5 M urea, 1% TFA, pH = 2.0, 4 °C | | | | | | | | | | | | | |
| Back-exchange | estimated from input recovery estimate of 0.7 and deuterated buffer concentration of 0.8 | | | | | | | | | | | | | |
| # of peptides | 87 | | | | | | | | | | | | | |
| Sequence coverage | 98% | | | | | | | | | | | | | |
| Average peptide length / Redundancy | 8 / 4.073 | | | | | | | | | | | | | |
| Replicates (biological or technical) | 3 (technical) | | | | | | | | | | | | | |
| Repeatability (average STD) | 3.542 | | | | | | | | | | | | | |
| Significant differences in HDX | > 5% D (unpaired t-tests at each time point, p-value < 0.01) | | | | | | | | | | | | | |
| **Data Set (Figure 6E, S15Bi)** | **WT Mpro** | | | | | | | **WT Mpro + NMTV** | | | | | | |
| Reaction details | 50 mM HEPES, 500 mM NaCl, 1 mM TCEP, pH 8.0, 4 °C | | | | | | | | | | | | | |
|  | 30 min | | | | | | | | | | | | | |
| Deuterium labeling time (sec) | 43200 | | | | | | | | | | | | | |
| HDX quench reaction details | 5 M urea, 1% TFA, pH = 2.0, 4 °C | | | | | | | | | | | | | |
| Back-exchange | estimated from input recovery estimate of 0.7 and deuterated buffer concentration of 0.8 | | | | | | | | | | | | | |
| # of peptides | 90 | | | | | | | | | | | | | |
| Sequence coverage | 98% | | | | | | | | | | | | | |
| Average peptide length / Redundancy | 8 / 4.163 | | | | | | | | | | | | | |
| Replicates (biological or technical) | 3 (technical) | | | | | | | | | | | | | |
| Repeatability (average STD) | 1.594 | | | | | | | | | | | | | |
| Significant differences in HDX | > 5 %D (unpaired t-tests at each time point, p-value < 0.01) | | | | | | | | | | | | | |
| **Data Set (Figure S15Bii)** | **WT Mpro** | | | | | | | **WT Mpro + Climbazole** | | | | | | |
| Reaction details | 50 mM HEPES, 500 mM NaCl, 1 mM TCEP, pH 8.0, 4 °C | | | | | | | | | | | | | |
|  | 30 min | | | | | | | | | | | | | |
| Deuterium labeling time (sec) | 43200 | | | | | | | | | | | | | |
| HDX quench reaction details | 5 M urea, 1% TFA, pH = 2.0, 4 °C | | | | | | | | | | | | | |
| Back-exchange | estimated from input recovery estimate of 0.7 and deuterated buffer concentration of 0.8 | | | | | | | | | | | | | |
| # of peptides | 90 | | | | | | | | | | | | | |
| Sequence coverage | 98% | | | | | | | | | | | | | |
| Average peptide length / Redundancy | 8 / 4.163 | | | | | | | | | | | | | |
| Replicates (biological or technical) | 3 (technical) | | | | | | | | | | | | | |
| Repeatability (average STD) | 1.808 | | | | | | | | | | | | | |
| Significant differences in HDX | > 5% D (unpaired t-tests at each time point, p-value < 0.01) | | | | | | | | | | | | | |
| **Data Set (Figure S15Bii)** | **WT Mpro** | | | | | | | **WT Mpro + Pelitinib** | | | | | | |
| Reaction details | 50 mM HEPES, 500 mM NaCl, 1 mM TCEP, pH 8.0, 4 °C | | | | | | | | | | | | | |
|  | 30 min | | | | | | | | | | | | | |
| Deuterium labeling time (sec) | 43200 | | | | | | | | | | | | | |
| HDX quench reaction details | 5 M urea, 1% TFA, pH = 2.0, 4 °C | | | | | | | | | | | | | |
| Back-exchange | estimated from input recovery estimate of 0.7 and deuterated buffer concentration of 0.8 | | | | | | | | | | | | | |
| # of peptides | 90 | | | | | | | | | | | | | |
| Sequence coverage | 98% | | | | | | | | | | | | | |
| Average peptide length / Redundancy | 8 / 4.163 | | | | | | | | | | | | | |
| Replicates (biological or technical) | 3 (technical) | | | | | | | | | | | | | |
| Repeatability (average STD) | 2.001 | | | | | | | | | | | | | |
| Significant differences in HDX | > 5% D (unpaired t-tests at each time point, p-value < 0.01) | | | | | | | | | | | | | |

Table S2. SEC-MALS-SAXS table for nsp7-11 and nsp7-8 polyprotein.

| A) **SAS data-collection parameters** | | | | |
| --- | --- | --- | --- | --- |
| Instrument | BioCAT facility at the Advanced Photon Source beamline 18ID with Pilatus3 X 1M (Dectris) detector | | | |
| Wavelength (Å) | 1.033 | | | |
| Beam size (μm^2^) | 150 (h) x 25 (v) focused at the detector | | | |
| Camera length (m) | 3.69 m for nsp7-11 sample and 3.631 m for the nsp7-8 sample | | | |
| q-measurement range (Å^-1^) | 0.003 to 0.35 | | | |
| Absolute scaling method | Glassy Carbon, NIST SRM 3600 | | | |
| Basis for normalization to constant counts | To transmitted intensity by beam-stop counter | | | |
| Method for monitoring radiation damage | Automated frame-by-frame comparison of relevant regions using CORMAP (5) implemented in BioXTAS RAW | | | |
| Exposure time, number of exposures | 0.5 s exposure time with a 1 s total exposure period (0.5 s on, 0.5 s off) of entire SEC elution | | | |
| Sample configuration | SEC-MALS-SAXS. Size separation used a Superdex 200 Increase 10/300 GL column and a 1260 Infinity II HPLC (Agilent Technologies). UV data was measured in the Agilent, and MALS-DLS-RI data by DAWN HELEOS-II (17 MALS + 1 DLS channels) and Optilab T-rEX (5) instruments (Wyatt Technology). SAXS data were measured in a sheath-flow cell (6), effective path length 0.542 mm | | | |
| Sample temperature (ºC) | 23 | | | |
| B) **Software employed for SAS data reduction, analysis and interpretation** | | | | |
| SAXS data reduction | Radial averaging; frame comparison, averaging, and subtraction done using BioXTAS RAW 2.0.3 (7) | | | |
| Basic analysis: Guinier, M.W., P(r) | Guinier fit and M.W. using BioXTAS RAW, P(r) function using GNOM (8). RAW uses MoW and Vc M.W. methods (4*,* 9) | | | |
| Shape/bead modelling | *DAMMIF* (*10*) and *DAMMIN* (*11*) via *ATSAS* | | | |
| MALS-DLS-RI analysis | Astra 7 (Wyatt) | | | |
| C) **Structural parameters** | | | | |
|  | **nsp7-11** | | **nsp-8** | |
|  | **monomer** | **dimer** | **monomer** | **dimer** |
| **Guinier analysis** |  | | | |
| I(0) | 2.98 +/- 4.18e-3 | 0.53 +/- 2.73e-3 | 0.79 +/- 2.61e-3 | 0.99 +/- 3.21e-3 |
| R_g_ (Å) | 34.91 +/- 0.15 | 45.69 +/- 0.39 | 24.96 +/- 0.16 | 33.02 +/- 0.2 |
| q_min_ (Å^-1^) | 0.00467 | 0.00297 | 0.00475 | 0.00475 |
| qR_g_max | 0.92 | 1.29 | 1.22 | 1.24 |
| Coefficient of correlation, R^2^ | 0.99 | 0.75 | 0.97 | 0.97 |
| M.W. (Bayes) [kDa] | 62.4 | 91.2 | 33.1 | 62.4 |
| **P(r) analysis** |  | | | |
| I(0) | 2.99 +/- 4.28e-3 | 0.53 +/- 3.59e-3 | 0.79 +/- 2.47e-3 | 0.99 +/- 3.58e-3 |
| R_g_ (Å) | 35.89 +/- 0.15 | 48.16 +/- 0.61 | 25.24 +/- 0.13 | 33.69 +/- 0.24 |
| d_max_ (Å) | 156.0 | 191.0 | 88.0 | 118.0 |
| q range (Å^-1^) | 0.005- 0.229 | 0.003- 0.175 | 0.005- 0.299 | 0.0047- 0.2424 |
| χ^2^ (total estimate from GNOM) | 2.277 (0.719) | 1.822 (0.771) | 1.085 (0.829) | 1.785 (0.793) |
| M.W. (Vp) [kDa] | 72.7 | 130.8 | 34.4 | 70.9 |
| Porod volume (Å^3^) | 8.75e+4 | 1.58e+5 | 4.15e+4 | 8.54e+4 |
| D) **Shape model-fitting results** | | | | |
|  | **nsp7-11 (monomer)** | | **nsp7-8 (monomer)** | |
| ***DAMMIF* (default parameters, 20 calculations)** |  | | | |
| q range for fitting (Å^-1^) | 0.005- 0.229 | | 0.005- 0.299 | |
| Symmetry, anisotropy assumptions | P1, none | | P1, none | |
| NSD (standard deviation), No. of clusters | 0.951 (0.057), 9 | | 1.087 (0.180), 7 | |
| χ^2^ range | 2.303-2.318 | | 1.092-1.093 | |
| Resolution (from SASRES) (Å) | 42 +/- 3 Å | | 27 +/- 2 Å | |
| M.W. estimate (kDa) | 80.4 | | 39.8 | |
| *DAMMIN* (default parameters) |  | | | |
| q range for fitting (Å^-1^) | 0.005- 0.229 | | 0.005- 0.299 | |
| Symmetry, anisotropy assumptions | P1, none | | P1, none | |
| χ^2^ | 2.278 | | 1.087 | |

**Table S3.** Summary of agreement of top ten nsp7-11 models with experimental data from HDX-MS, XL-MS, SAXS, and junction accessible surface area.

|  | | Model A1 | Model A2 | Model A3 | Model B1 | Model B2 | Model B3 | Model B4 | Model C1 | Model C2 | Model D1 |
| --- | --- | --- | --- | --- | --- | --- | --- | --- | --- | --- | --- |
| RMSE  from HDXer | | 0.2402 | 0.2568 | 0.2264 | 0.2522 | 0.2849 | 0.2657 | 0.2886 | 0.3092 | 0.3108 | 0.2864 |
| % XL satisfied  (≤ 30 Å) | | 75.3 | 76.3 | 75.3 | 85.6 | 92.8 | 89.7 | 91.8 | 88.7 | 92.8 | 78.4 |
| Rg (Å) | | 32.60 | 30.99 | 32.37 | 29.45 | 28.77 | 28.77 | 28.57 | 27.26 | 27.78 | 29.52 |
| 𝛘^2^ | | 6.42 | 15.53 | 7.89 | 20.76 | 29.12 | 31.48 | 34.55 | 49.0 | 40.91 | 18.31 |
| Junction Accessible Surface Area (Å^2^) | **nsp7-8 site** | 414.7 | 425.9 | 393.4 | 155.7 | 161.3 | 160.0 | 155.5 | 142.5 | 182.3 | 141.6 |
|  | **nsp8-9 site** | 280.6 | 276.3 | 201.9 | 353.2 | 138.7 | 251.4 | 209.8 | 97.3 | 129.9 | 208.6 |
|  | **nsp9-10 site** | 219.3 | 324.7 | 308.2 | 306.5 | 336.0 | 340.1 | 232.9 | 440.1 | 357.4 | 420.8 |
|  | **nsp10-11 site** | 248.2 | 254.1 | 270.6 | 292.1 | 396.1 | 273.4 | 223.2 | 247.0 | 310.1 | 261.5 |

**Table S4.** SARS-CoV-2 Mpro crystallographic binders interacting in various regions of Mpro showing protection from solvent exchange in differential HDX-MS of C145A Mpro versus C145A Mpro with nsp7-11 at 12 h incubation in deuterated buffer only.

| **PDB code** | **PDB chains** | **Ligand PDB code** | **Region of Mpro** | **Fragment or Small molecule (name)** | **Antiviral activity** | **Pubchem CID** | **PMID** |
| --- | --- | --- | --- | --- | --- | --- | --- |
| 5RE5 | A | T0J_A_404 | Back of the catalytic domain | Fragment | ND | 769265 | 33028810 |
| 5RE6 | A | O0S_A_404 |  | Fragment | ND | 1487531 | 33028810 |
| 5RFB | A | K3S_A_404 |  | Fragment | ND | 62755740 | 33028810 |
| 5RFC | A | K1Y_A_404 |  | Fragment | ND | 811874 | 33028810 |
| 5RGG | A | NZD_A_404 |  | Fragment | ND | 762797 | 33028810 |
| 5RH4 | A | UHG_A_1001 |  | Fragment | ND | 6997572 | 33028810 |
| **6YVF** | **A** | **A82_A_401** |  | **AZD6482** | **NA** | **44137675** | **33811162** |
| f | | | | | | | |
| 5REF | A | 6SU_A_404 | Near dimerization domain | Fragment | ND | 2806372 | 33028810 |
| **7AGA** | **A** | **LZE_A_401** |  | **AT7519** | **EC_50_ = 25 μM** | **11338033** | **33811162** |
| f | | | | | | | |
| 5REG | A | LWA_A_404 | Surface pocket | Fragment | ND | 1086839 | 33028810 |
| **7AOL** | **A** | **RQH_A_403** |  | **climbazole** | **NA** | **25271637** | **33811162** |
| f | | | | | | | |
| 5RFA | A | JGY_A_403 | Dimerization domain | Fragment | ND | 1224835 | 33028810 |
| **7ABU** | **A** | **R6Q_A_401** |  | **RS102895** | **EC_50_ = 19.8 μM** | **10000456** | **33811162** |
| **7AMJ** | **A** | **RMZ_A_409** |  | **PD 168568** | **NA** | **9798466** | **33811162** |
| **7AXM** | **A** | **93J_A_502** |  | **pelitinib** | **EC_50_ = 1.25 μM** | **6445562** | **33811162** |
| 7BFB | B | 9JT_B_1005 |  | Ebselen | EC_50_ = 4.67 μM* | 126410 | 32272481 |

ND = not determined; NA = no activity; ligands assayed in bold; *this small molecule binds in two more sites in Mpro; ligands in bold were tested in limited proteolysis assay and colored according to **Fig 6A**.
